# Differences in evolutionary accessibility determine which equally effective regulatory motif evolves to generate pulses

**DOI:** 10.1101/2020.12.02.409151

**Authors:** Kun Xiong, Mark Gerstein, Joanna Masel

**Affiliations:** Department of Molecular and Cellular Biology, University of Arizona, Tucson AZ, 85721; Department of Molecular Biophysics and Biochemistry, Yale University, New Haven CT, 06511; Program in Computational Biology and Bioinformatics, Yale University, New Haven CT, 06511; Department of Computer Science, Yale University, New Haven CT, 06511; Department of Statistics and Data Science, Yale University, New Haven CT, 06511; Department of Ecology and Evolutionary Biology, University of Arizona, Tucson AZ, 85721

**Keywords:** Pulse generation, adaptationism, mutation-biased adaptation, gene regulatory network, transcriptional regulation

## Abstract

Transcriptional regulatory networks (TRNs) are enriched for certain “motifs”. Motif usage is commonly interpreted in adaptationist terms, i.e. that the optimal motif evolves. But certain motifs can also evolve more easily than others. Here, we computationally evolved TRNs to produce a pulse of an effector protein. Two well-known motifs, type 1 incoherent feed-forward loops (I1FFLs) and negative feedback loops (NFBLs), evolved as the primary solutions. Which motif evolves more often depends on selection conditions, but under all conditions, either motif achieves similar performance. I1FFLs generally evolve more often than NFBLs, unless we select for a tall pulse. I1FFLs are more evolutionarily accessible early on, before the effector protein evolves high expression; when NFBLs subsequently evolve, they tend to do so from a conjugated I1FFL-NFBL genotype. In the empirical *S. cerevisiae* TRN, output genes of NFBLs had higher expression levels than those of I1FFLs. These results suggest that evolutionary accessibility, and not relative functionality, shapes which motifs evolve in TRNs, and does so as a function of the expression levels of particular genes.

## INTRODUCTION

The topology of transcriptional regulatory networks (TRNs) is enriched for certain motifs (Lee et al. 2002; Milo et al. 2002; Shen-Orr et al. 2002; Mangan and Alon 2003). Many argue that these motifs are the result of adaptive evolution where the motif whose dynamical behavior best provides the beneficial function is the one that will evolve (Alon 2007). However, adaptationist claims about TRN organization have been accused of being just-so stories, with adaptive hypotheses still in need of testing against an appropriate null model of network evolution (Wagner 2003; Artzy-Randrup et al. 2004; Mazurie et al. 2005; Kuo et al. 2006; Solé and Valverde 2006; Lynch 2007; Knabe et al. 2008; Jenkins and Stekel 2010; Tsuda and Kawata 2010; Widder et al. 2012; Ruths and Nakhleh 2013; Payne and Wagner 2015). We recently generated such a null model and used it to show that coherent type 1 feed-forward loops can, as hypothesized, evolve adaptively in response to selection to filter out short spurious signals, by combining a fast signaling pathway and a slow signaling pathway with an AND gate (Xiong et al. 2019). Testing the hypothesis in this way was not merely confirmatory, but generated other insights about the existence and nature of alternative adaptive solutions, especially when slow transcriptional regulation is combined with faster response mechanisms such as post-translational regulation (Xiong et al. 2019). Other network motifs and properties have not yet received similar treatment.

At least three different motifs (**Fig. 1A**) are all capable of producing a sharp pulse of expression in response to an increase in input signal (**Fig. 1B**) (Basu et al. 2004; Camas et al. 2006; Çağatay et al. 2009). All depend on an effector first being rapidly activated by a signal, and later, at a slower timescale, being repressed by it. These three motifs are simple auto-repression (AR), negative feed-back loops (NFBLs), and incoherent type 1 feed-forward loops (I1FFLs) (**Fig. 1A**). The three motifs are topologically and functionally similar to each other, differing in whether the slow repression is effected via negative autoregulation by the effector *R* of itself, via negative feedback regulation of *R* using a specialized repressor, or via a separate negative control pathway from the input to the repressor and then the effector.

**Figure 1.**
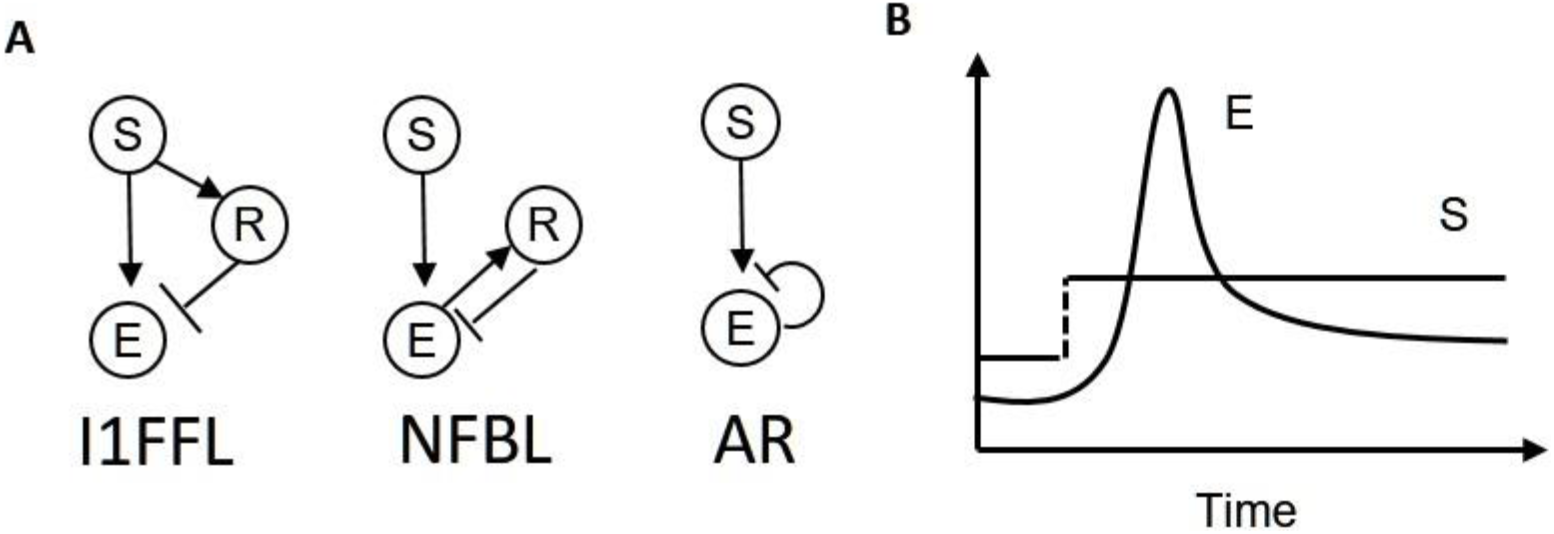
Three motifs (I1FFL, NFBL, and AR) all produce a pulse of effector E expression in response to increased signal S. **(A)** In all three cases, rapid and direct activation of the effector by the signal is eventually countered by a slower path of repression. The three motifs differ topologically in whether repression is by the effector itself (AR), by a specialized repressor (R) that is activated by the signal (I1FFL), or by a specialized repressor that is activated by the effector (NFBL). Regular arrow tips represent activation and ⊣ represents repression. **(B)** With appropriate parameters, and with a delay between transcriptional activation and protein production in the case of AR, all three motifs can induce a pulse, as the initial increase in expression as S activates E is eventually tamped down by a path of repression.

The high prevalence of I1FFLs and NFBLs in TRNs has been interpreted to occur because these two motifs are adaptations for pulse generation and closely related functions (Shoval and Alon 2010; Shoval et al. 2010; Ferrell 2016; Shi et al. 2017). Both I1FFLs and NFBLs allow the steady-state level of the effector, before and after the pulse, to be independent of the signal strength, a property known as chemical adaptation (Ferrell 2016; Shi et al. 2017). We note that AR is normally hypothesized to perform functions other than pulse generation (Wall et al. 2004; Alon 2007), but theoretical analysis and experiments show that AR can generate pulses (Rosenfeld et al. 2002; Camas et al. 2006; Amit et al. 2007). We therefore include AR for the completeness of the study, while focusing on I1FFLs and NFBLs.

Which of the motifs is likely to evolve is often explained by adaptive demands for specific properties of the pulse. For example, although both I1FFLs and NFBLs allow the amplitude of the pulse to be a function of the fold-change of the signal’s strength (Shoval et al. 2010), they do so with different functional forms (Adler et al. 2014). I1FFLs and NFBLs can also differ in their ability to filter noise in the signal (Buzi and Khammash 2016).

Alternatively, non-adaptive causes might be responsible for differences in the occurrence of the three motifs. An important non-adaptive consideration is that fitness landscapes tend to have many alternative local endpoints, which might take the form either of peaks (Whitlock et al. 1995) or of plateaus (van Nimwegen and Crutchfield 2000). Factors such as expression levels can change the relative accessibility of different local evolutionary endpoints, in ways that are independent of differences in their heights. Note that by “non-adaptive” explanations, we do not mean “neutral evolution”. Instead we refer to evolutionary accessibility, encompassing both which mutations occur in a single step and which hill-climbing multi-step paths are possible. This emphasis on *process* as non-adaptive explanation is in contrast to adaptive explanations that consider only the optimality of the final evolutionary *outcome*. Whether the non-adaptive explanation of evolutionary accessibility is a plausible cause is a question that *in silico* evolution is ideally set up to explore. We note that I1FFLs and NFBLs differ by whether it is the signal or the effector that regulates the repressor (**Fig. 1A**). Intuitively, the relative ease of evolving these two possible regulatory interactions with the repressor could depend on the relative expression levels of the candidate regulators.

Here we simulate TRN evolution under selection to produce a pulse, and test how subtle differences between scenarios might have both adaptive and non-adaptive effects on which motifs evolves. In particular, a highly expressed effector is more able to stimulate its repressor, and we therefore predict that this scenario should be more likely to evolve regulation via an NFBL and correspondingly less likely to evolve an I1FFL. Our simulations reject adaptationist explanations – I1FFLs and NFBLs achieve similar fitness – and confirm that NFBLs are evolutionarily more accessible than I1FFLs under this scenario, but that I1FFLs are more accessible under other scenarios where a highly expressed effector is not required. Data from real-world yeast TRNs agree with model predictions, showing that the effectors of NFBLs generally have higher expression levels than those of I1FFLs.

## MATERIALS AND METHODS

### Transcriptional regulation

Transcription factors (TFs) bind to a given TF binding site (TFBS) according to a formula based on the biophysics of the matching of the cis-regulatory sequence to the TF’s consensus binding sequence (see Supplementary Materials/TF binding for details). Briefly, in isolation from all other TFs and TFBSs, the probability *P_b_* that a TFBS is occupied is

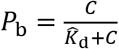

where *C* is the total concentration of the TF and 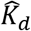 is a version of the binding affinity *K_d_* of the TF, rescaled to account for the fact that focal TFBSs must compete for TF with many non-specific binding sites throughout the genome. From probabilities of this form, we calculate the probability that exactly A activators and R repressors are bound to a given cis-regulatory sequence, given the possibility of physical overlap among TFBSs (see Supplementary Materials/TF occupancy). From this, we derive four probabilities that we assume regulate gene expression: 1) the probability *P_A_* of having at least one activator bound to a gene, 2) the probability *P_R_* of having at least one repressor bound, 3) the probability *P*_A_no_R_ of having no repressors and one activators bound, and the probability *P*_notA_no_R_ of having no TFs bound.

We model transcriptional initiation as a two-step process whose rates depend on TF binding, and parameterize those rates with reference to nucleosome disassembly followed by transcription machinery assembly (Mao et al. 2010; Brown et al. 2013). We model a repressed state of nucleosome presence, an intermediate state of a nucleosome-free transcription start site that lacks transcription machinery, and an active state. We set the transition rate from the repressed state to the active state to

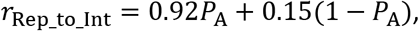

using as bounds for our linear function 0.15 min^−1^ as the background rate of histone acetylation (Katan-Khaykovich and Struhl 2002) (which leads to nucleosome disassembly) and 0.92 min^−1^ as the rate of nucleosome disassembly for the constitutively active PHO5 promoter (Brown et al. 2013).

We set the transition rate from the intermediate state to the active state to

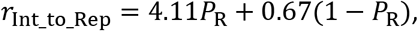

where 0.67 min^−1^ is a background histone de-acetylation rate (Katan-Khaykovich and Struhl 2002) and 4.11 min^−1^ is chosen so as to keep a similar maximum:basal rate ratio as that of *r*_Rep_to_Int_.

We assume that the binding of a single repressor can prevent the transition from the intermediate state to the active state (Courey and Jia 2001). In the absence of repressors, activators facilitate the assembly of transcription machinery (Poss et al. 2013). Under these assumptions, we set the transition rate from the intermediate state to the active state to

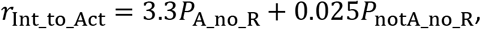

where 3.3 min^−1^ is the rate of transcription machinery assembly for a constitutively active *PHO5* promoter (Brown et al. 2013), and 0.025 min^−1^ is same rate when the *PHO4* activator of the *PHO5* promoter is knocked out.

We set the transition rate *r*_Act_to_Int_ from the active state to the intermediate state to be gene-specific and independent of TF binding. This is because the promoter sequence not only determines which specific TFBSs are present, but also influences non-specific components of the transcriptional machinery (Decker and Hinton 2013). See Supplementary materials/*r*_Act_to_Int_ for the parameterization of *r*_Act_to_Int_.

### Fitness

Our simulations of gene expression begin with a burn-in phase of random length, to ensure that TRNs to respond to a change in the signal, rather than evolve a timer mechanism. The level of signal is low during the stage one burn-in, which lasts for 120 + x minutes, where x is random number drawn from an exponential distribution truncated at 30, and with an un-truncated average of 10. Fitness is assessed only on the basis of stage two, which lasts for 240 minutes, plus the last 5 minutes of stage one. We sample the effector concentration at one-minute intervals. The highest effector concentration during stage two is denoted *p*.

The fitness of a TRN has four components: the peak level of effector, a low effector expression starting point, the speed with which effector expression rises, and the speed with which it falls. Together, these four components capture the core attributes of what it means to be a pulse, and in combination, they apply consistent selective pressure first to generate any pulse at all and later to produce a superior pulse. All four fitness components are based on the expression level of the effector. For the purpose of scoring effector concentration and hence fitness, we use the total protein level of all effector proteins, including those that have diverged, following duplication, to have different regulatory activities.

Fitness component one scores the match to a pre-defined peak effector expression level:

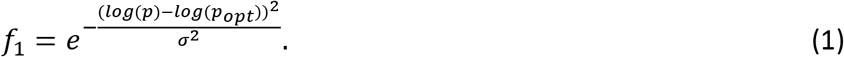

We set the optimal peak expression level *p_opt_* to 5,000 molecules per cell, 10,000 molecules per cell, or 20,000 molecules per cell, corresponding to selection for a low, medium, or high peak level, respectively. Under the assumption that the effector is a metabolism-related protein, we chose the number 10,000 based on the average number of PDC1 protein molecules per yeast cell (Ghaemmaghami et al. 2003). The effector also acts as TF; this kind of dual functionality is not uncommon in yeast (Gancedo and Flores 2008). We set *σ*^2^ = 0.693 so that when *p* = 0.5*p_opt_*, *f_1_* = 0.5.

We set fitness component two to reward low effector expression at the end of stage one:

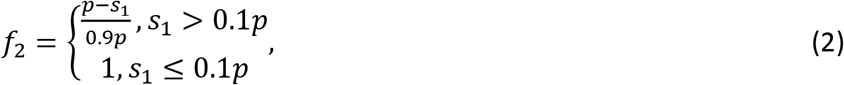

where *s_1_* is the arithmetic mean of the effector level across the last 5 minutes of stage one. This is chosen as a simple piecewise-linear function, which plateaus at a maximum of 1 for values of *s_1_* below 10% of the peak level *p*.

We set fitness component three to reward rapid turn-on of effector:

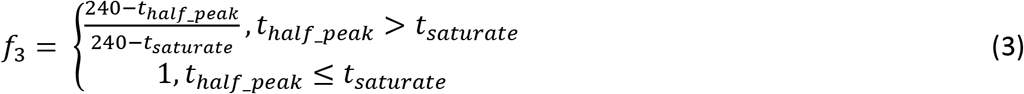

where *t_half_peak_* is the latest time in stage 2 at which the effector level is at 0.5(*s_1_* + *p*) before the effector hits its peak, and *t_saturate_* sets a time for which making effector response still more rapid no longer increases fitness. We set *t_saturate_* to 60 minutes.

To select for the downward slope of a pulse, fitness component four rewards falls in the effector falls to no more than 80% of the peak level by the end of stage two:

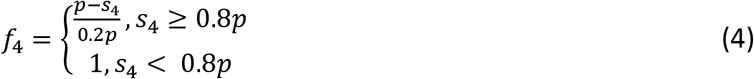

where *s_4_* is the arithmetic mean of the effector level across the last 5 minutes of stage two. Again, we chose a piecewise-linear function. We chose the relatively high value of 80% in order to select for an inclusive category of pulses. We consider pulses that eventually return all the way down to the level that prevailed before the signal (i.e. biochemical adaptation) to be a special case.

In some simulations of gene expression, we observed a peak expression level that is smaller than or equal to the effector’s expression level right before the signal increases, or is even 0. In neither of these cases do fitness components Eq. 2 and/or 4 provide a useful selection gradient toward the evolution of a pulse. For simplicity, we set the fitness of these two cases to zero.

In addition to the selection described above to favor a pulse, at each point in the simulation, gene expression also incurs a cost that is proportional to the total rate of translation of all genes (see Supplementary Materials/Cost of gene expression). The estimated fitness of a TRN from one gene expression simulation is the arithmetic mean of the four components minus the cumulative cost of gene expression throughout the last 360 minutes of a simulation of gene expression.

### Evolution

We calculate the arithmetic mean fitness *f_resident_* of the current (“resident”) TRN across 1000 replicate simulations of gene expression, and the arithmetic mean fitness *f_mutant_* of the mutant across 200 replicate simulations of gene expression. If *f_mutant_* satisfies

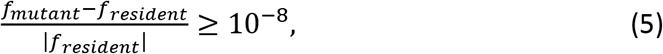

we replace the resident TRN with the mutant, and re-calculate the fitness of the new resident TRN to higher resolution using an additional 800 replicate simulations of gene expression.

Because gene expression is stochastic in our simulations, estimated fitness varies among replicates, and is subject to error even after averaging across many replicates. This means that our algorithm allows neutral or slightly deleterious mutations to fix. This is sometimes even explicit; the updated fitness that includes 800 additional simulations of the successful mutant can be lower than the fitness of the TRN it replaced.

Standard origin-fixation evolutionary simulations explicitly calculate a probability of fixation for each mutation and compare it to a pseudo-random number to decide whether fixation occurs. Our model achieves a similar exploration of nearly neutral evolutionary paths by using the intrinsic uncertainty in the stochastic estimation of fitness. Our approach wastes as few beneficial mutations as possible, minimizing computation, rather than discard most beneficial mutations through the use of a fixation probability that is only around twice the selection coefficient (Haldane 1927). For example, in our simulations, we accepted 0.5 million out of 1.9 million trialed mutations across 10 evolution replicates in the high-peak condition, of which only a minority can be presumed to have achieved true fitness increases (**Fig. S1**). Importantly, fixation probability in our algorithm still depends on the size of the true underlying fitness difference, which controls the probability that the estimated selection coefficient in Eq. 5 will be positive.

### Counting network motifs

We count I1FFLs and NFBLs formed by the signal, an effector gene, and a repressor gene that is different from the effector gene, with interactions between them as shown in **Fig. 1A**. We count ARs formed by the signal and an effector gene. We score gene A as potentially regulating gene B, i.e. creating one of the links shown in **Fig. 1A**, if there is a TFBS for A in the cis-regulatory sequence of B. We allow genes in I1FFLs and NFBLs to self-regulate. An overlapping I1FFL in which the effector and the auxiliary TF repress each other is counted not as two I1FFLs, but rather as a different (and rarer) type of network motif. Overlapping I1FFLs evolve rarely.

Given that two mismatches to an 8-bp consensus sequence still yield above-background binding, a random 8-bp sequence qualifies as a weak affinity TFBS with probability 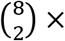 0.75^2^ × 0.25^6^ = 0.0038. Each cis-regulatory sequence contains around 300 8-bp potential binding sites (including both orientations of a 150 bp cis-regulatory sequence), among which 1.14 will on average qualify by chance as a two-mismatch TFBS for a given TF. These two-mismatch TFBSs, occurring so often by chance, usually have low affinity, and therefore might have little regulatory effect. It is for this reason we refer to them above as potential regulatory interactions – our previous work has shown that motifs can appear more clearly when weak affinity TFBSs with little regulatory effect are excluded (Xiong et al. 2019). Four types of spurious two-mismatch TFBSs can create apparent but non-functional I1FFLs and NFBLs: S → TF, E → TF, TF → E, and E → E (**Fig. S2**), where “TF” refers here to a transcription factor that is not an effector. Because it is computationally expensive to test whether each two-mismatch TFBS is spurious, we instead tested all cases at a time for each of the four types listed above. Specifically, we recalculate the fitness of the TRN while ignoring all 2-mismatch TFBSs of that type, across 1,000 gene expression simulations, and deem the entire set of TFBSs spurious if the recalculated fitness is at least 99% of the original fitness (see **Fig. 2** legend for variations on this criterion). We ignore spurious connections while scoring network motifs.

**Figure 2.**
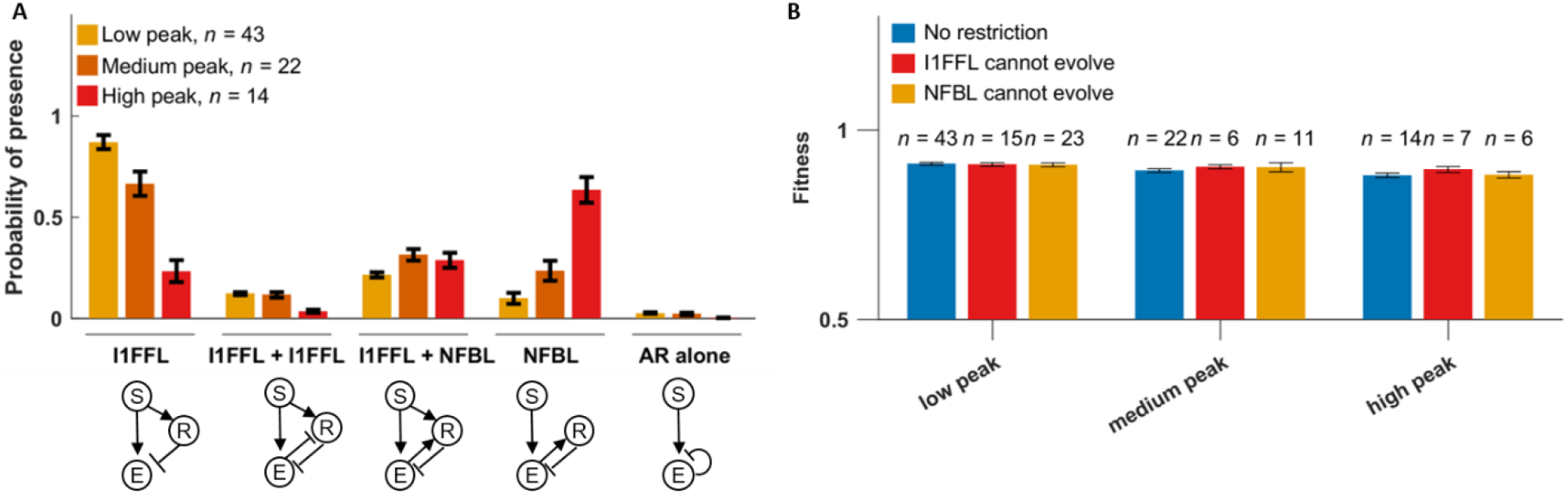
Selection for high peak effector expression levels promotes NFBLs. **(A)** TRNs are evolved under selection to generate pulses in response to an input signal. Under all three selection conditions, the input signal starts with 100 molecules per cell and increases to 1,000 molecules per cell to trigger a pulse. Three versions of the evolutionary simulations select for three different optimal peak effector levels of the effector: low (*p_opt_* = 5,000 molecules per cell), medium (*p_opt_* = 10,000 molecules per cell), and high (*p_opt_* = 20,000 molecules per cell). For each high-fitness genotype (**Fig. S4**), we calculate the proportion of evolutionary steps that contain at least one network motif of the specified type among the last 10,000 evolutionary steps (out of a total of 50,000 evolutionary steps). When scoring for motifs, non-functional spurious TFBSs were excluded (see Methods for details, and **Fig. S6** for results using different TFBSs exclusion criteria). R can be auto-regulating (not shown in circuit diagram). On rare occasions, AR co-occurred with I1FFLs or overlapping I1FFLs (labelled here I1FFL + I1FFL) (**Fig. S7**), and these few cases were included in the scoring of I1FFL and overlapping I1FFL frequencies. **(B)** Preventing either NFBLs or I1FFLs from evolving does not lower the final fitness within high-fitness evolutionary simulations. Instead, genotypes obtained equally high fitness by evolving the other common motif (**Fig. S8**). To prevent NFBLs from evolving, we remove the TF binding activity of effectors; this also prevents the evolution of the AR auto-repression motif. To prevent I1FFLs from evolving, we ignore TFBSs for the signal in the cis-regulatory sequence of any repressors. Because this might have unintended consequences for mutations that convert repressors to/from activators, we set to zero the rate of mutations that effect this conversion. Data are shown as mean ± SE over replicates.

### Mutations that create and destroy motifs

For each evolutionary replicate, we identified the evolutionary steps at which the number of instances of a given motif changes to or from zero, which we call “motif-destroying-mutations” and “motif-creating-mutations”, respectively. We removed spurious TFBSs before scoring motifs, as described in the last section, with one modification: to save computation related to mutations that were trialed and then rejected by selection, we used only 200 gene expression simulations to determine fitness without the TFBS in question, with a threshold of 98% of original fitness. Mutations that change the expression levels of a gene and/or the binding affinity of a TF can potentially change whether a two-mismatch TFBS is “spurious” in terms of fitness effects, effectively rewiring the TRNs even if they do not create or destroy core TFBSs of the motif in question.

### Expression levels of TFs in yeast TRN

We used YeastMine (Balakrishnan et al. 2012) to retrieve 129 *S. cerevisiae* genes that have the GO term “DNA-binding transcription factor activity” or children of this GO term. We then searched Yeastract (Teixeira et al. 2006) for TFs that regulate these 129 TFs, demanding evidence from both DNA binding and gene expression. When the search found new TFs that are not included in list given by YeastMine, the new TFs were added to the list and fed to Yeastract again. We stopped the iterative search when no new TFs were found, and the final list has 203 TFs. Yeastract annotates interactions between pairs of TFs as activating, repressing, or both. When annotated as “both” (i.e. likely condition-specific), we interpreted it as whichever interaction mode would be needed in order to complete a motif. We scored I1FFLs, NFBLs, and their conjugates from all combinations of three TFs out of the 203, allowing E and/or R to be self-repressing. Because the effectors of NFBLs must be activators, we excluded I1FFLs whose effectors are repressors in case there is a systematic difference in expression between activators and repressors. In total, we identified 46 NFBLs, 30 I1FFLs, and 7 I1FFL-NFBL conjugates.

To assess peak expression level, we used the data of Gasch et al. (2000), who applied multiple stimuli to yeast and measured the fold-change in RNA expression of all genes relative to pre-stimulus expression levels. We analyzed data on exposure to 10 stimuli: amino acid starvation, nitrogen depletion, sorbitol osmotic shock, temperature shift from 25° to 37°, diamide, hydrogen peroxide, menadione disulfate, diauxic shift, dithiothreitol, and transition to a stationary phase of growth. Following each stimulus, fold-change was recorded over several time points. We consider an effector gene to exhibit pulse-like expression if the maximum fold-increase in expression occurs prior to the last time point and has a larger magnitude than that of the maximum fold-decrease in expression; we excluded gene-stimulus combinations that do not meet this criterion from further analysis. For input and repressor genes, we did not require a pulse, but merely that the stimulus led to increased expression (measured as average fold-change across time points), and that the maximum fold-increase was larger than the maximum fold-decrease. We excluded repressor-stimulus and input-stimulus combinations that failed to meet both criteria.

We note that the same gene can occupy the same position within multiple motifs. For example, GAT1 is the effector in 18 NFBLs and one I1FFL, suggesting that this gene might be particularly well-suited for function within NFBLs. To compare gene expression between I1FFLs and NFBLs, we weighted the fold-change in expression of a given gene by the frequency with which that gene appears in the motif of interest, e.g. weights of 18/19 and 1/19 for GAT1’s appearance as an effector in NFBLs and I1FFLs, respectively. For I1FFL-NFBL conjugates, we assign half-weights to both I1FFLs and NFBLs.

We complemented this peak-RNA-expression analysis with an analysis of the average protein levels (i.e. not peak levels), taken from PaxDB (Wang and Purisima 2005). One analysis is restricted to a March 2013 data set originally compiled by PeptideAtlas (Desiere et al. 2006) to show the abundances of peptides in *S. cerevisiae* pooled across 90 experiments, which include normal growth conditions and perturbed growth conditions, e.g. cell cycle arrest and metabolic perturbation. We also used another data set “GPM, Aug, 2014” from PaxDB, which has more genes than “PeptideAtlas, March 2013” (5289 versus 4828). While we could not find a detailed description for this GPM (the Global Proteome Machine) (Craig et al. 2004) dataset, GPM generally includes data from PeptideAtlas (Craig et al. 2004), meaning that this data similarly includes both normal growth conditions and perturbed growth conditions. Weighted average protein levels were calculated with the same weighting scheme as for fold-change of gene expression.

### Data Availability

The source code for our computational model is available at https://github.com/MaselLab/network-evolution-simulator/tree/I1_FFLs.

## RESULTS

### Model overview

We used a previously described computational model to simulate the expression of genes in a TRN, parameterized by available *Saccharomyces cerevisiae* data (Xiong et al. 2019). **Fig. S3** summarizes the model, and the model parameters are summarized in **Tables S1** and **S2**. The TRN evolves under a realistic mutational spectrum including *de novo* appearance of weak-affinity TFBSs, and frequent gene duplication and deletion. Briefly, each gene in the TRN encodes either an activating or repressing TF, and each is regulated by a 150-bp cis-regulatory sequence accessible to TF binding. Each TF recognizes an 8-bp consensus binding sequence with a characteristic binding affinity. Binding sites with up to two mismatches are still recognized, with each mismatch reducing binding affinity according to a thermodynamic model (Supplementary Materials/TF Binding). TFs can bind in either orientation. Each TF that binds to DNA occupies three extra base pairs upstream and downstream of the consensus sequence, making a total of 14 bp inaccessible to other TFs. The concentrations of TFs are used to calculate the probabilities that each cis-regulatory region is bound by a given number of activators and repressors (see Methods).

To simulate gene expression, we assume that each gene transitions between an active chromatin state that can initiate transcription, an intermediate primed state capable of becoming either activated or repressed, and a repressed chromatin state. Most transition rates depend on whether activators and/or repressors are bound (see Methods), with the fastest transition rate to the active state occurring when at least one activator and no repressors are bound. The transcription initiation rates of mRNAs from active genes are gene-specific, and so are the degradation rates. Note that the above rates (including the transition rates between the states of genes) are expectations; exactly when a reaction (e.g. one of gene A’s mRNAs is degraded) happens is simulated stochastically using a Gillespie algorithm (Gillespie 1977). Conceptually, the algorithm allows one event to happen at a time, with the cellular state remaining unchanged between events. The waiting time between two events has an exponential distribution, with a mean specified by the total reaction rates. Once the time of an event is sampled, the algorithm randomly picks an event (e.g. degrading gene A’s mRNA) based on the reaction’s relative rate, and changes the cellular state according to the event (e.g. there is one less mRNA of gene A in cell). See Supplementary materials for details.

Each mRNA produces protein at a gene-specific translation rate. Once transcription is initiated, we simulate a delay before mRNA can be translated at full speed. The delay accounts for the completion of both transcription and the loading of ribosomes to mRNA, and is a function of gene length (Supplementary materials/Transcriptional delay and Translational delay). Because tracking the turnover of individual protein molecules with a Gillespie algorithm is computationally expensive, we calculate the turnover of proteins with ordinary differential equations (Supplementary materials/Simulation of gene expression).

To select for pulse generation, we designate an input signal to the TRN, which binds to cis-regulatory regions like any other TF, but whose concentration is set externally rather than being regulated by other TFs in the TRN. The input signal always activates gene expression. Signal concentration is low and constant during a burn-in phase, where genes are initialized with a repressed chromatin state, and begin with zero non-signal mRNA and protein. Then in stage 2, the signal instantly switches to a high level, and selection is applied for a TF designated to be the “effector” to exhibit pulse-like expression. High fitness depends on having low effector expression at the end of stage 1, matching a pre-defined peak effector concentration during stage 2, rapidly increasing effector level after stage 2 begins, and having a low effector level at end of stage 2. Details of the signal and fitness calculation are given in the Methods.

We initialize an evolutionary simulation with a randomly generated genotype of 3 activator genes, 3 repressor genes, and an effector gene. The effector is initialized as an activator, which makes NFBLs more accessible than ARs (although below we will explore the effects of switching this). All quantitative gene-specific parameter values, such as transcriptional rates and gene length, are randomly initialized according to empirically estimated distributions (see **Table S1** and Supplementary materials).

We simulate five classes of mutations. **Table S2** lists the corresponding mutation rates and details of the parameterization are provided in the Supplementary materials. A class-one mutation is a duplication or deletion of one gene along with its cis-regulatory sequence. The maximum number of genes is capped at four effector genes plus 21 non-effector genes (excluding the signal) to limit computational cost. Once this limit is reached, no duplication mutations are allowed. In addition, once any give gene is present in four copies, none of the copies are duplicated until one is again lost by deletion. Neither the last effector gene nor the last non-effector gene are subject to deletion. The signal is subject neither to duplication nor to deletion.

Class-two mutations are single nucleotide substitutions in the cis-regulatory sequences, which can cause TFBS turnover. Mutations change one nucleotide to one of the other three nucleotides with equal probabilities.

Class-three mutations change quantitative gene-specific parameters, i.e. the rate at which transcriptional bursts end, gene length, mRNA degradation rate, protein synthesis rate, protein degradation rate, and the affinity of a TF to DNA. All quantitative gene-specific parameters except length are subjected to mutational bias, e.g. mutation tends to reduce the affinity of TF binding. In case this is insufficient to ensure the values of the mutatable parameters never go beyond reasonable limits, we also apply hard bounds (see Supplementary materials/Mutations for details).

Class-four mutations convert transcription activators to repressors (or the reverse). This mutation does not apply to the input signal, i.e. the input signal is always an activator.

Class-five mutations change a single nucleotide preference in a TF’s consensus binding sequence. One of the other three nucleotides is chosen for the consensus binding sequence with equal probabilities.

When gene duplicates differ due only to class-three mutations, the duplicates are considered as “copies” of the same gene, encoding “protein variants”. Once a class-four or class-five mutation is applied to a gene duplicate, the duplicate becomes a new gene encoding a new protein. When scoring motifs, we require that each node be a different protein.

Evolution is simulated using the revised origin-fixation model introduced by Xiong et al. (2019). Briefly, the resident genotype experiences one mutation, chosen according to the relative rates of all possible mutations. The fitness of the original resident TRN and of the mutant TRN is calculated by simulating gene expression in response to an input signal (see Methods for details). If the estimated fitness of the mutant is sufficiently high (see Methods for details), the mutant replaces the resident genotype. Note that estimated fitnesses include stochasticity from the simulation of gene expression, which serves to introduce a form of genetic drift. If no replacement occurs, we generate a new mutant and repeat the procedure until a replacement is found. We call a replacement an evolutionary step, and end each simulation after 50,000 evolutionary steps. We use the average fitness of the last 10,000 evolutionary steps to determine whether evolution has found a good solution.

### High peak expression level non-adaptively promotes NFBLs

We evolve TRNs under selection to generate a pulse of effector expression in response to a sudden 10-fold increase in input. While any of the three network motifs can solve this challenge, a highly expressed effector is more capable of stimulating its repressor, and thus this solution should be more likely to evolve regulation via an NFBL and correspondingly less likely to evolve an I1FFL. Note that this prediction is expected on both adaptive grounds of which solution might be superior, and on non-adaptive grounds of which solution is easier for evolution to find.

To test this prediction, we vary the optimal peak level of the effector (see Methods for details of fitness function). *In silico* evolution from a random starting point is not always successful at reaching the target phenotype, so we focus on the most evolutionarily successful simulations. We do this by dividing evolutionary replicates into three categories based on final fitness (**Fig. S4**). See **Fig. S5** for examples of the phenotypes of the high-fitness replicates.

High-fitness solutions rarely involve AR under any of the three selection conditions, while both I1FFLs and NFBLs evolve often (**Fig. 2A**). As predicted, when we select for higher effector expression, we get more NFBLs and fewer I1FFLs (**Fig. 2A**). These NFBLs were absent from medium-fitness solutions, which instead employed I1FFLs or ARs (**Fig. S9A**), generally achieving lower peak effector expression than in the high-fitness solutions (**Fig. S9B**). While this seems to suggest that NFBLs might be adaptively superior, if we prevent one type of motif from evolving, similarly high fitness genotypes can be obtained via the other motif (**Figs. 2B and S8**). The reason we get more NFBLs and fewer I1FFLs with selection for higher peak effector expression is therefore not straightforward adaptive superiority of the former, but rather the relative ease of finding high-fitness solutions.

### Early bias toward I1FFLs can shift to later NFBL evolution via I1FFL-NFBL conjugates

The combined frequency of the two motifs rises throughout the long period of evolution, rather than topological solutions being found early and becoming locked in and only incrementally improved on. However, the frequency of I1FFLs in particular rises prominently during the first 10,000 evolutionary steps (**Fig. 3**), even under selection for high peak effector levels, i.e. selection that ultimately leads to an evolutionary preference for NFBLs (**Fig. 3C**).

**Figure 3.**
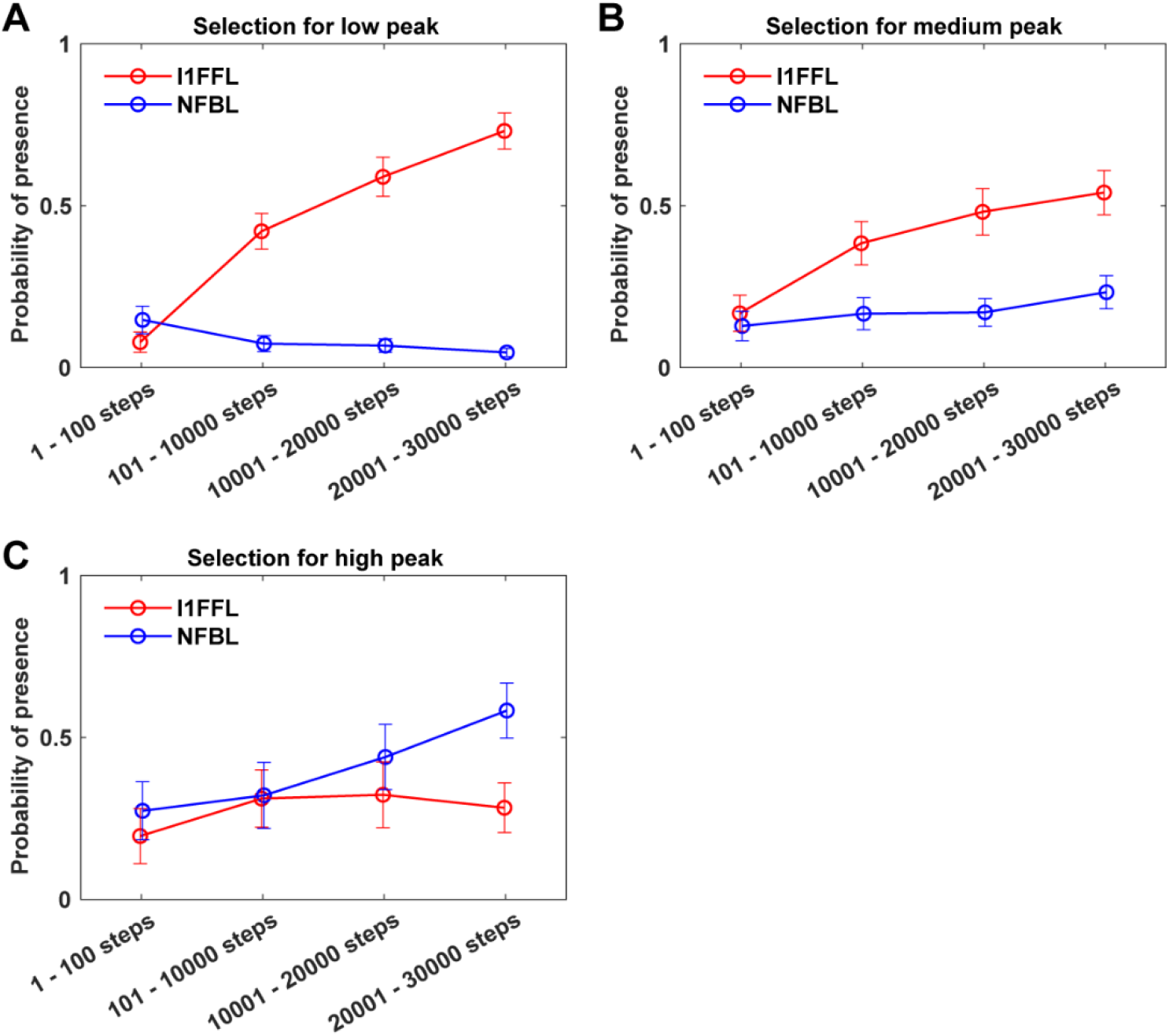
Evolution of I1FFLs and NFBLs follow different trajectories. We score motif occurrence during different time periods along the way to the evolution of the high-fitness replicates shown in **Fig. 2A**. See **Fig. S10** for the occurrence of other motifs during evolution. As in **Fig. 2A**, we calculated the proportion of evolutionary steps that contain at least one network motif of the specified type. Note that because some evolutionary replicates oscillate between motif presence and absence, given the potential for slightly deleterious mutations in our evolutionary algorithm, the fraction of evolutionary replicates that frequently show the motif in question is higher than the probability of presence in one evolutionary step as shown here. Data are shown as mean ± SE over replicates.

To further test this point, we made the early evolution of NFBLs less accessible by initializing the effector as a repressor. While this reduced the frequency of NFBLs even under selection for a high peak, those NFBLs that still evolved reached similar performance to I1FFLs (**Fig. S11**). This further supports early evolutionary accessibility as a key factor.

The relative ease of I1FFL evolution could be because more mutations create I1FFLs and/or because mutations creating I1FFLs have higher acceptance rates. To explore this further, we characterize the mutations that create I1FFLs and/or NFBLs in TRNs that do not currently contain such a motif. I1FFL-creating mutations occur at a higher rate than NFBL-creating mutations under selection for low-peak and medium-peak expression, while NFBL-creating mutations are more common under selection for high-peak expression **(Table 1)**. The rarity of NFBL-creating mutations becomes much more pronounced when we restrict our analysis to mutations that do not also destroy or create another motif – this tendency holds even under conditions that favor NFBLs, i.e. late in evolution under selection for high-peak expression (**Table 1**). Greater mutational accessibility of the I1FFL motif is clearly one of the factors favoring this motif.

**Table 1.** Summary of mutations that create I1FFLs and/or NFBLs. We identify the accepted and rejected mutations that increase the number of I1FFLs and/or NFBLs in a TRN to above zero (see Methods for details). Among these mutations, “non-disruptive mutations” are those that create the given motif but do not otherwise alter the numbers of I1FFL (when NFBLs are created), NFBLs (when I1FFLs are created), I1FFL-NFBL conjugates, overlapping I1FFLs, and auto-repressors. For each selection condition and evolutionary stage, we pooled the qualified mutations from all high-fitness replicates shown in **Fig. 2A**. The total numbers of mutations of the given type were normalized by dividing by the total number of mutations trialed in resident TRNs that did not already have the motif in question. The acceptance rate shown in the table is the number of accepted mutations across all replicates divided by the number of trialed mutations across all replicates. Pseudoreplication may be a concern here; if the initial TRN tends to create one motif over the other, this might be propagated at all subsequent time points for that evolutionary replicate. However, **Table S3** shows that the initial mutational bias of a TRN can flip at a later stage of evolution.

| peak level |  | Evolutionary step 1 – 10,000 |  |  |  | Evolutionary step 10,001 – 30,000 |  |  |  |
| --- | --- | --- | --- | --- | --- | --- | --- | --- | --- |
|  |  | all mutations |  | non-disruptive mutations |  | all mutations |  | non-disruptive mutations |  |
|  |  | Trialed | Acceptance rate | Trialed | Acceptance rate | Trialed | Acceptance rate | Trialed | Acceptance rate |
| Low | I1FFL-creating | 0.049 | 0.171 | 0.0038 | 0.606 | 0.077 | 0.142 | 0.0019 | 0.550 |
|  | NFBL-creating | 0.026 | 0.078 | 0.00074 | 0.544 | 0.024 | 0.062 | 0.00028 | 0.504 |
| Medium | I1FFL-creating | 0.072 | 0.102 | 0.0031 | 0.607 | 0.088 | 0.083 | 0.00091 | 0.432 |
|  | NFBL-creating | 0.043 | 0.097 | 0.00084 | 0.694 | 0.049 | 0.078 | 0.00032 | 0.507 |
| High | I1FFL-creating | 0.036 | 0.114 | 0.0017 | 0.546 | 0.039 | 0.063 | 0.00027 | 0.411 |
|  | NFBL-creating | 0.097 | 0.072 | 0.0017 | 0.642 | 0.147 | 0.069 | 0.00026 | 0.476 |

The early evolution of I1FFLs is also facilitated by the higher acceptance rate of I1FFL-creating mutations relative to NFBL-creating mutations, particularly during the first 10,000 evolutionary steps (**Table 1)**. Similarly, I1FFL-destroying mutations are accepted less often than NFBL-destroying mutations are, in this case throughout the course of evolution and regardless of target peak expression (**Table S4**). Note that mutations that create one motif frequently destroy another, with NFBL-creating mutations more prone to this problem than I1FFL-creating mutations (**Table 2**). While some such disruptive mutations are accepted by our evolutionary algorithm (**Table 2**), acceptance rates are higher for non-disruptive mutations (**Table 1**). If we restrict our analysis to non-disruptive mutations, we see stronger mutation bias toward I1FFLs, and more similar acceptance rates for I1FFLs vs NFBLs (**Table 1**). In other words, a shortage of non-disruptive NFBL-creating mutations is an obstacle to the evolution of NFBLs. NFBL-creating mutations that destroy I1FFL-NFBL conjugates are both more common and more likely to be accepted than NFBL-creating mutations that destroy I1FFLs (**Table 2**). This suggests that I1FFL-NFBL conjugates might be an important intermediate step in the evolution of NFBLs, rather than NFBLs evolving de novo. This makes sense; after early evolution of an I1FFL provides a partial solution to the selective challenge, the evolutionary path to an NFBL does not abandon that I1FFL solution, but instead passes through a combined I1FFL-NFBL intermediate. The evolutionary path from an early partial I1FFL solution might lead either to a superior I1FFL or to an NFBL, with the potential to achieve similarly high fitness in either case.

**Table 2.** Most NFBL-creating mutations also destroy other motifs. A high fraction of trialed mutations that create a given motif also destroy I1FFL-NFBL conjugates, and many also destroy NFBLs (in the case of I1FFL-creating mutations) or I1FFLs (in the case of NFBL-creating mutations). Destructive mutations are accepted at significant rates. Qualified mutations are pooled across all evolutionary replicates. See Methods for details about the identification of mutations that create and/or destroy motifs.

| peak level |  | Evolutionary step 1-10,000 |  |  |  | Evolutionary step 10,001-30,000 |  |  |  |
| --- | --- | --- | --- | --- | --- | --- | --- | --- | --- |
|  |  | Destroys I-N conjugates | accept. rate | Destroys I or N | accept. rate | Destroys I-N conjugates | accept. rate | Destroys I or N | accept. rate |
| Low | I1FFL-creating | 0.461 | 0.108 | 0.101 | 0.169 | 0.529 | 0.092 | 0.106 | 0.183 |
|  | NFBL-creating | 0.924 | 0.060 | 0.443 | 0.037 | 0.883 | 0.048 | 0.443 | 0.034 |
| Medium | I1FFL-creating | 0.643 | 0.048 | 0.100 | 0.137 | 0.647 | 0.056 | 0.132 | 0.108 |
|  | NFBL-creating | 0.920 | 0.082 | 0.314 | 0.051 | 0.928 | 0.070 | 0.280 | 0.053 |
| High | I1FFL-creating | 0.466 | 0.047 | 0.255 | 0.148 | 0.611 | 0.021 | 0.274 | 0.090 |
|  | NFBL-creating | 0.962 | 0.060 | 0.307 | 0.051 | 0.962 | 0.064 | 0.234 | 0.047 |

Indeed, I1FFL-NFBL conjugates (and NFBLs) are also often converted by mutation into simple I1FFLs. However, under selection for high peak effector expression, the acceptance rate of such mutations decreases over evolutionary time (**Table 2**). Peak effector expression increases during evolution (**Fig. S12**); this could drive increased preference for the now more highly expressed effector rather than the signal to control the repressor. In medium-fitness evolutionary replicates, high peak effector expression is not achieved, and NFBLs rarely evolve (**Fig. S9**). By the same logic, we hypothesize that strengthening the input signal should promote I1FFLs even under selection for a high effector peak. This is indeed the case, with promotion in particular of the evolution of the I1FFL-NFBL conjugate (**Fig. S13**).

### Highly expressed effectors tend to be regulated by NFBLs in yeast

Next we tested our model predictions about when I1FFLs vs. NFBLs tend to evolve. We identified NFBLs, I1FFLs and I1FFL-NFBL conjugates in the TRN of *S. cerevisiae*, using Yeastract annotations of regulatory interactions between TFs (see Methods). Using data from Gasch et al. (2000), we identified genes that display pulse-like expression in response to an environmental stimulus, and the peak heights of the pulses (measured as the fold-change of RNA expression levels relative to the expression level before the stimulus). In agreement with our model prediction, the effectors of NFBLs reach higher peaks than those of I1FFLs following stimulus (**Fig. 4A**). However, the input signals of NFBLs increase their expression more in response to stimuli than do those of I1FFLs (**Fig. 4A**), which disagrees with our model prediction. We note that the 46 NFBLs in our dataset involve 26 unique genes as the input signal and 8 as the effector, while the 30 I1FFLs involve 14 signals and 9 effectors. This raises the possibility that the more diverse signal inputs of the NFBLs might contain more false positive hits.

**Figure 4.**
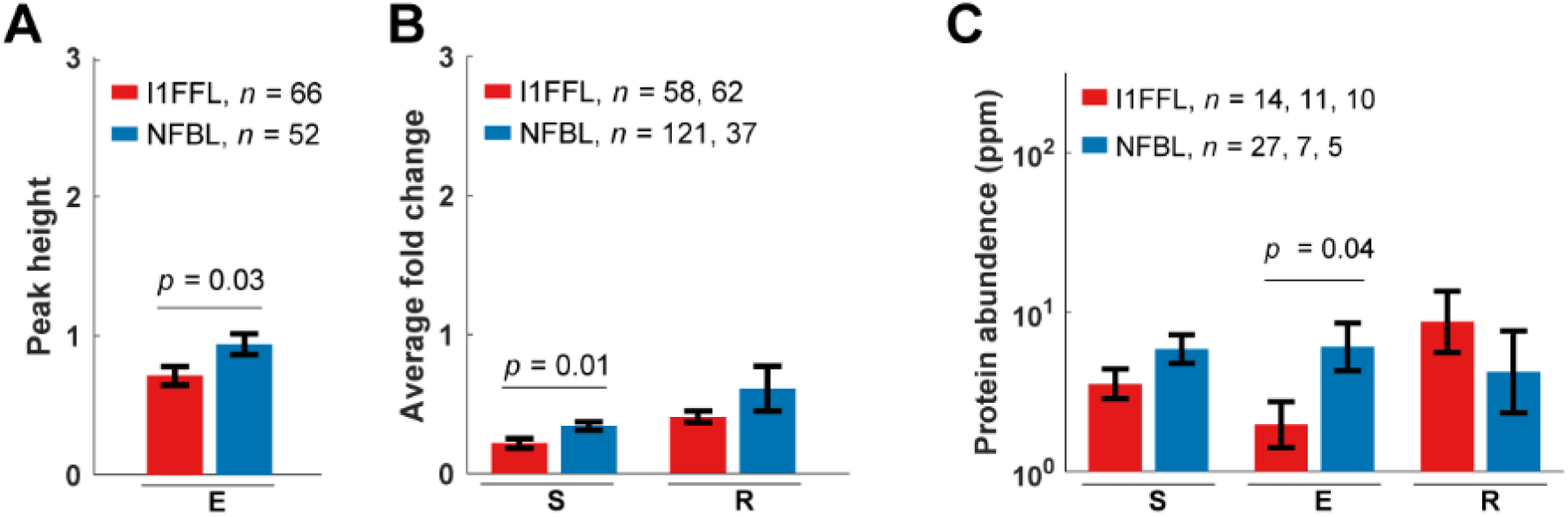
Effector TFs in yeast NFBLs have higher expression than those in I1FFLs. **(A)** Peak height of pulses was measured as the maximum fold-increase in RNA expression in response to one of 10 stimuli (see methods for details), for the subset of genes showing pulse-like RNA expression during the former in the RNA expression data of Gasch et al. (2000). **(B)** Average fold-change in signal and repressor RNA expression in response to stimuli, for the subset that showed an increase (see Methods). **(C)** Protein levels under normal conditions were taken from the “PeptideAtlas, March, 2013” dataset provided by PaxDB (Wang et al. 2015). A weaker result was obtained using a different dataset from PaxDB that includes a larger set of gene-environment combinations (**Fig. S14**). For fold-change in expression, data are shown as mean ± SE over each network position across all instances of the motif. The procedure is similar for protein abundance, except the data is first log transformed. For each motif, we list the numbers *n* of unique gene-stimulus combinations where pulse-like expression is observed at a signal node (S), effector node E, or repressor node R. p-values come from two-tailed t-tests.

We also analyzed yeast protein expression levels from PaxDB, averaged across multiple environmental conditions rather than measured in response to stimuli (see Methods). We found that effector TFs generally have higher expression in NFBLs than in I1FFLs (**Fig. 4B**). Note that the direction of causation is not known from the empirical data alone: when an effector already has high expression this might prompt the evolution of NFBL, or the presence of an NFBL might facilitate the evolution of high effector expression. The theoretical work presented here presents non-exclusive proof of principle in support of the former interpretation.

## DISCUSSION

We selected for a pulse generator in an evolutionary simulation model and observed which TRN motifs emerged. As predicted, selecting for high peak expression level of the effector promotes NFBLs over I1FFLs, while a strong input signal promotes I1FFLs. However, if one motif is prevented from evolving, the other motif can evolve to take its place, with no loss of peak fitness, suggesting that the preference between motifs is not adaptive in origin, i.e. is not about which motif is optimal. One predicted pattern is confirmed in the actual TRN of *S. cerevisiae*, where the effector’s expression level is higher in NFBLs than in I1FFLs.

Both mutational accessibility (i.e. how often mutations create the given motif) and selective acceptance rates (Yampolsky and Stoltzfus 2001; Stoltzfus and McCandlish 2017; Gomez et al. 2020) contribute to patterns of relative evolutionary accessibility. Note that the motif created by larger-effect beneficial mutations need not be better at generating a pulse. The latter is what is meant by an “adaptive” explanation for the dominance of I1FFL over NFBL (or the vice versa) (Gould and Lewontin 1979). A non-adaptive evolutionary explanation can include a role of selection or an increase in fitness during evolution, but emphasizes process rather than final fitness as the cause of bias in evolutionary outcomes.

Usually mutational accessibility and selective acceptance rates point in the same direction, but not always: I1FFLs are less mutationally accessible under early selection for high peak effector expression, but have a relatively high mutation acceptance rate. The higher acceptance rates for I1FFL-creating mutations do not reflect functional superiority of I1FFLs, but rather the fact that creating NFBLs frequently involves destroying other, likely functional, motifs. Avoidance of damage to existing functions has been previously noted in other discussions of the evolutionary paths taken by TRNs (Wagner 2003; Carroll 2008; Stern and Orgogozo 2009; Sorrells and Johnson 2015). The mutational accessibility of different motifs is not static, but changes over the course of an evolutionary path (**Table S3**).

We did not pit the performance of I1FFLs and NFBLs versus the evolutionary accessibility of the two motifs, because both motifs had indistinguishable performance in our system. We are therefore unable to answer whether the evolutionary accessibility of motifs can alter the evolutionary outcome predicted by performance of motifs. However, finding in our system that high fitness solutions can often be found one way or another is intriguing in its possible generality.

We find that most NFBLs evolve not from connecting previously disconnected genes (e.g. S->E->R), but rather from uncoupling I1FFL-NFBL conjugates in favor of a pure NFBL. We simulate only relatively small TRNs, due to limitations in computational power, and this might restrict the evolutionary trajectories that are capable of generating network motifs. If simulation algorithms that scaled better with TRN size were devised, it would be interesting to explore whether network motifs would evolve via different trajectories in larger TRNs. For example, the use of the same TF for multiple regulatory purposes in real-world TRNs, which of course are larger, can constrain network evolution, requiring complex trajectories to achieve a new regulatory function (Sorrells et al. 2015).

We predicted via simulations that a highly expressed effector should promote the evolution of NFBLs over I1FFLs. Strikingly, this prediction was borne out in empirical data from yeast. A highly expressed TF can more strongly regulate its target, and/or reduce the amount of noise propagated downstream (Pedraza and van Oudenaarden 2005; Jothi et al. 2009). Once a highly expressed TF gains a TFBS in the target gene, the TFBS may also be easier to retain during evolution. Many studies on TRNs have noted a systematic difference among the expression levels of genes at topologically different positions (Herrgård et al. 2003; Yu et al. 2003; Jothi et al. 2009; Gerstein et al. 2012), and that highly expressed TFs are often regulators of multiple target genes (Jothi et al. 2009; Gerstein et al. 2012). Our findings also support the idea that the observed network motifs in TRNs are partially shaped by the expression levels of TFs.

## Acknowledgement

We acknowledge the generous computational resources provided by the High Performance Computation Center of the University of Arizona.

## Supplementary materials

**Table S1.**
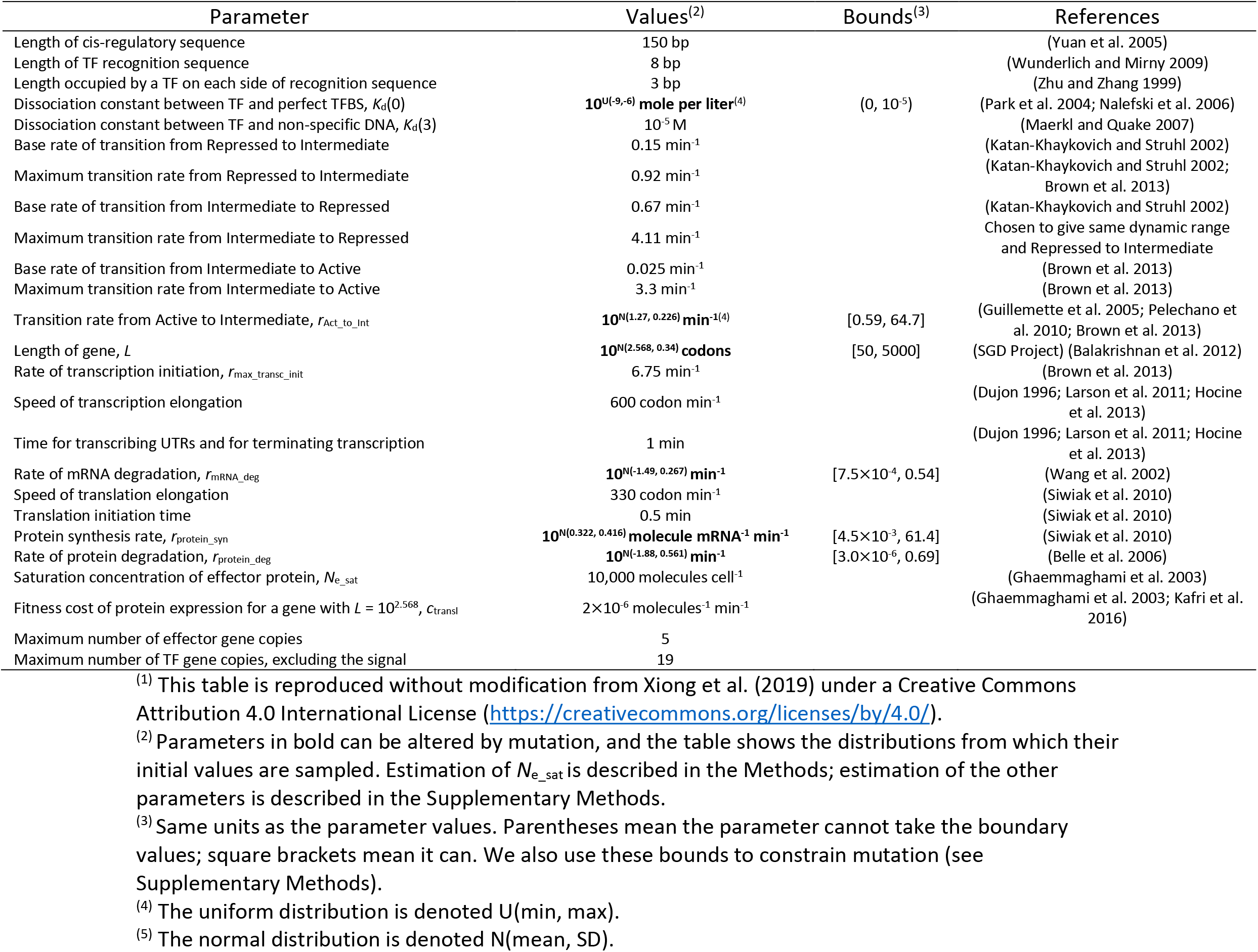
Major model parameters^(1)^

**Table S2.**
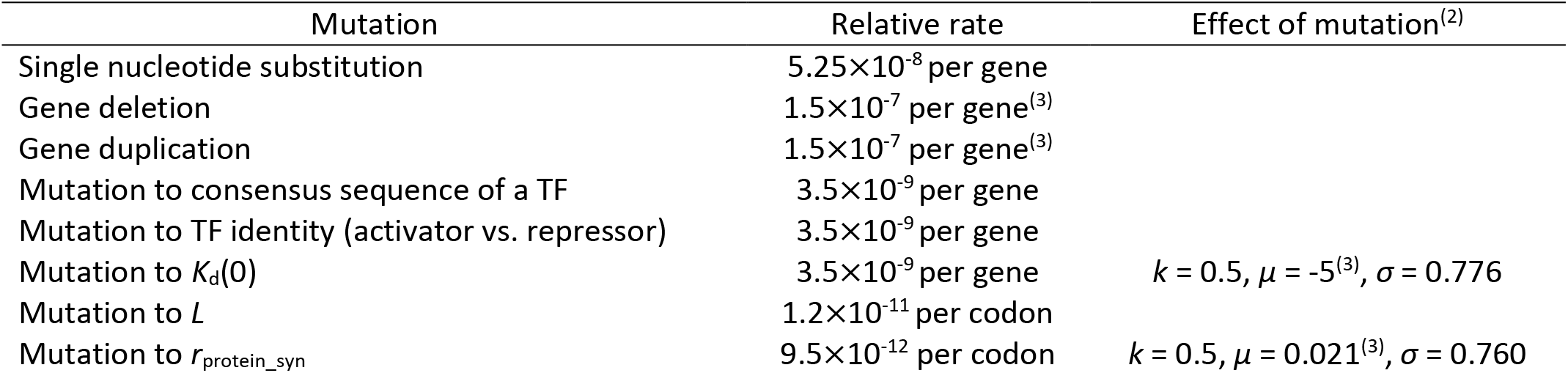
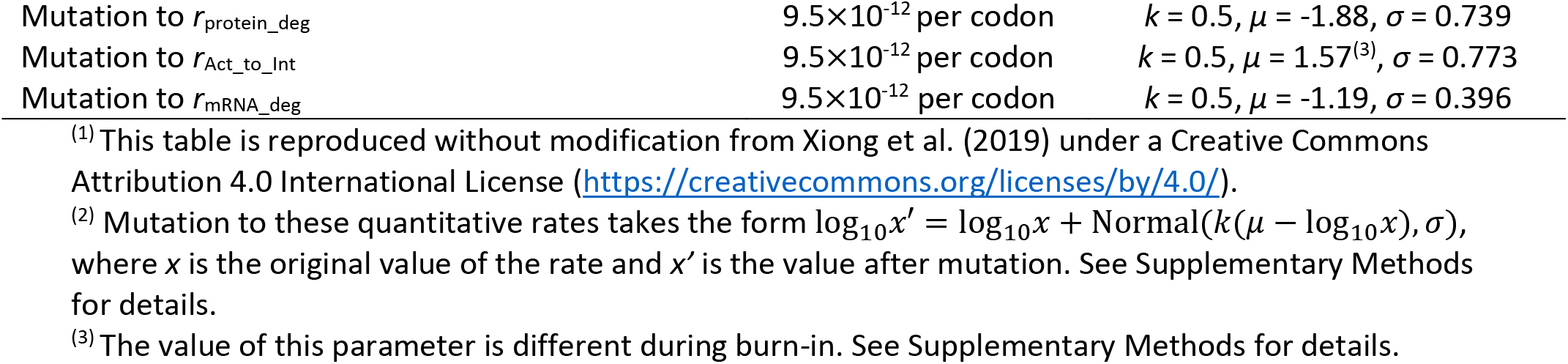
Mutation rates and effect sizes^(1)^

| Mutation | Relative rate | Effect of mutation<sup>(2)</sup> |
| --- | --- | --- |
| Single nucleotide substitution | $5.25 \times 10^{-8}$ per gene | |
| Gene deletion | $1.5 \times 10^{-7}$ per gene<sup>(3)</sup> | |
| Gene duplication | $1.5 \times 10^{-7}$ per gene<sup>(3)</sup> | |
| Mutation to consensus sequence of a TF | $3.5 \times 10^{-9}$ per gene | |
| Mutation to TF identity (activator vs. repressor) | $3.5 \times 10^{-9}$ per gene | |
| Mutation to $K_d(0)$ | $3.5 \times 10^{-9}$ per gene | $k = 0.5, \mu = -5^{(3)}, \sigma = 0.776$ |
| Mutation to $L$ | $1.2 \times 10^{-11}$ per codon | |
| Mutation to $r_{protein\_syn}$ | $9.5 \times 10^{-12}$ per codon | $k = 0.5, \mu = 0.021^{(3)}, \sigma = 0.760$ |
| Mutation to $r_{\text{protein\_deg}}$ | $9.5 \times 10^{-12}$ per codon | $k = 0.5, \mu = -1.88, \sigma = 0.739$ |
| Mutation to $r_{\text{Act\_to\_Int}}$ | $9.5 \times 10^{-12}$ per codon | $k = 0.5, \mu = 1.57^{(3)}, \sigma = 0.773$ |
| Mutation to $r_{\text{mRNA\_deg}}$ | $9.5 \times 10^{-12}$ per codon | $k = 0.5, \mu = -1.19, \sigma = 0.396$ |

<sup>(1)</sup> This table is reproduced without modification from Xiong et al. (2019) under a Creative Commons Attribution 4.0 International License (https://creativecommons.org/licenses/by/4.0/).
<sup>(2)</sup> Mutation to these quantitative rates takes the form $\log_{10} x' = \log_{10} x + \text{Normal}(k(\mu - \log_{10} x), \sigma)$ , where $x$ is the original value of the rate and $x'$ is the value after mutation. See Supplementary Methods for details.
<sup>(3)</sup> The value of this parameter is different during burn-in. See Supplementary Methods for details.

**Table S3.** Mutational bias toward particular motifs can shift over the course of evolution. We focus our analysis on three random TRN initializations (conditions 25, 31, and 79) that evolved to high fitness in all three selection conditions. Under selection for high peak effector expression, all three simulations evolved NFBLs (i.e. the occurrence of NFBL > 0.5 and the occurrence of I1FFL < 0.5). Under selection for low or medium effector expression, all three evolved I1FFLs. As in Table 1, we show the number of mutations normalized by the total number of mutations trialed in resident TRNs that did not contain the motif in question. As an example of a change in mutational bias, initial condition 25 under selection for low peak effector expression initially creates NFBLs more often but later creates I1FFLs more often.

| Optimal peak |  | Initial condition 25 |  | Initial condition 31 |  | Initial condition 79 |  |
| --- | --- | --- | --- | --- | --- | --- | --- |
|  |  | Evo. step 1 - 10,000 | Evo. step 10,001 – 20,000 | Evo. step 1 - 5,000 | Evo. step 5,001 – 10,000 | Evo. step 1 - 10,000 | Evo. step 10,001 – 20,000 |
| Low | I1FFL-creating | 0.050 | 0.217 | 0.680 | 0.067 | 0.051 | 0.586 |
|  | NFBL-creating | 0.107 | 0.007 | 0.006 | 0.003 | 0.106 | 0.013 |
| Medium | I1FFL-creating | 0.065 | 0.075 | 0.811 | 0.365 | 0.105 | 0.047 |
|  | NFBL-creating | 0.111 | 0.104 | 0.016 | 0.025 | 0.112 | 0.171 |
| High | I1FFL-creating | 0.013 | 0.034 | 0.875 | 0.059 | 0.048 | 0.118 |
|  | NFBL-creating | 0.729 | 0.413 | 0.007 | 0.201 | 0.259 | 0.174 |

**Table S4.**
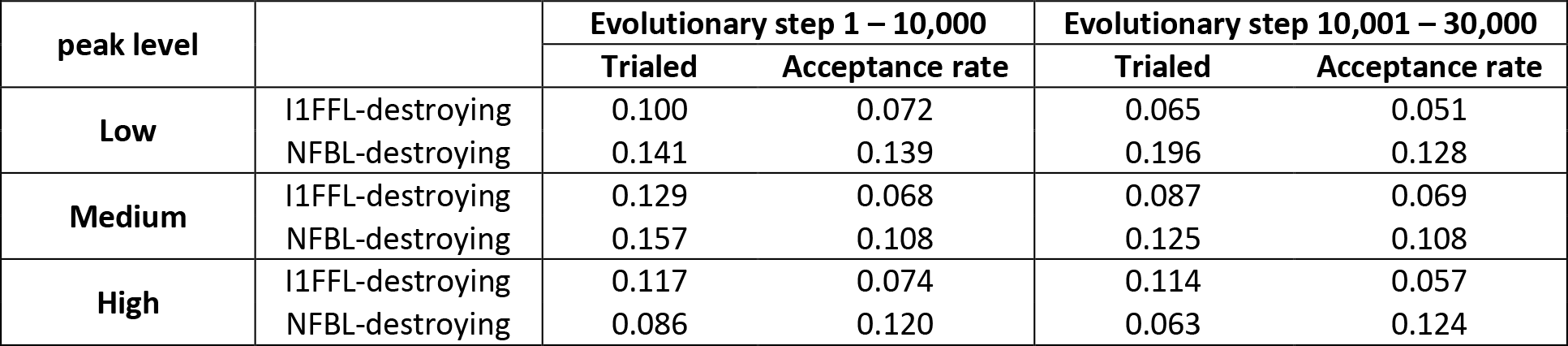
Summary of mutations that remove all I1FFLs and/or NFBLs. For each selection condition, we pooled qualified mutations from all high-fitness replicates shown in Fig. 2. A mutation is classed as destroying if it eliminates all instances of the given motif. The total number of qualified mutations were normalized by the total number of mutations trialed in resident TRNs that contained the motif of interest. The acceptance rate is the number of accepted mutations across all replicates divided by the number of trialed mutations across all replicates.

| peak level |  | Evolutionary step 1 – 10,000 |  | Evolutionary step 10,001 – 30,000 |  |
| --- | --- | --- | --- | --- | --- |
|  |  | Trialed | Acceptance rate | Trialed | Acceptance rate |
| Low | I1FFL-destroying | 0.100 | 0.072 | 0.065 | 0.051 |
|  | NFBL-destroying | 0.141 | 0.139 | 0.196 | 0.128 |
| Medium | I1FFL-destroying | 0.129 | 0.068 | 0.087 | 0.069 |
|  | NFBL-destroying | 0.157 | 0.108 | 0.125 | 0.108 |
| High | I1FFL-destroying | 0.117 | 0.074 | 0.114 | 0.057 |
|  | NFBL-destroying | 0.086 | 0.120 | 0.063 | 0.124 |

**Figure S1.**
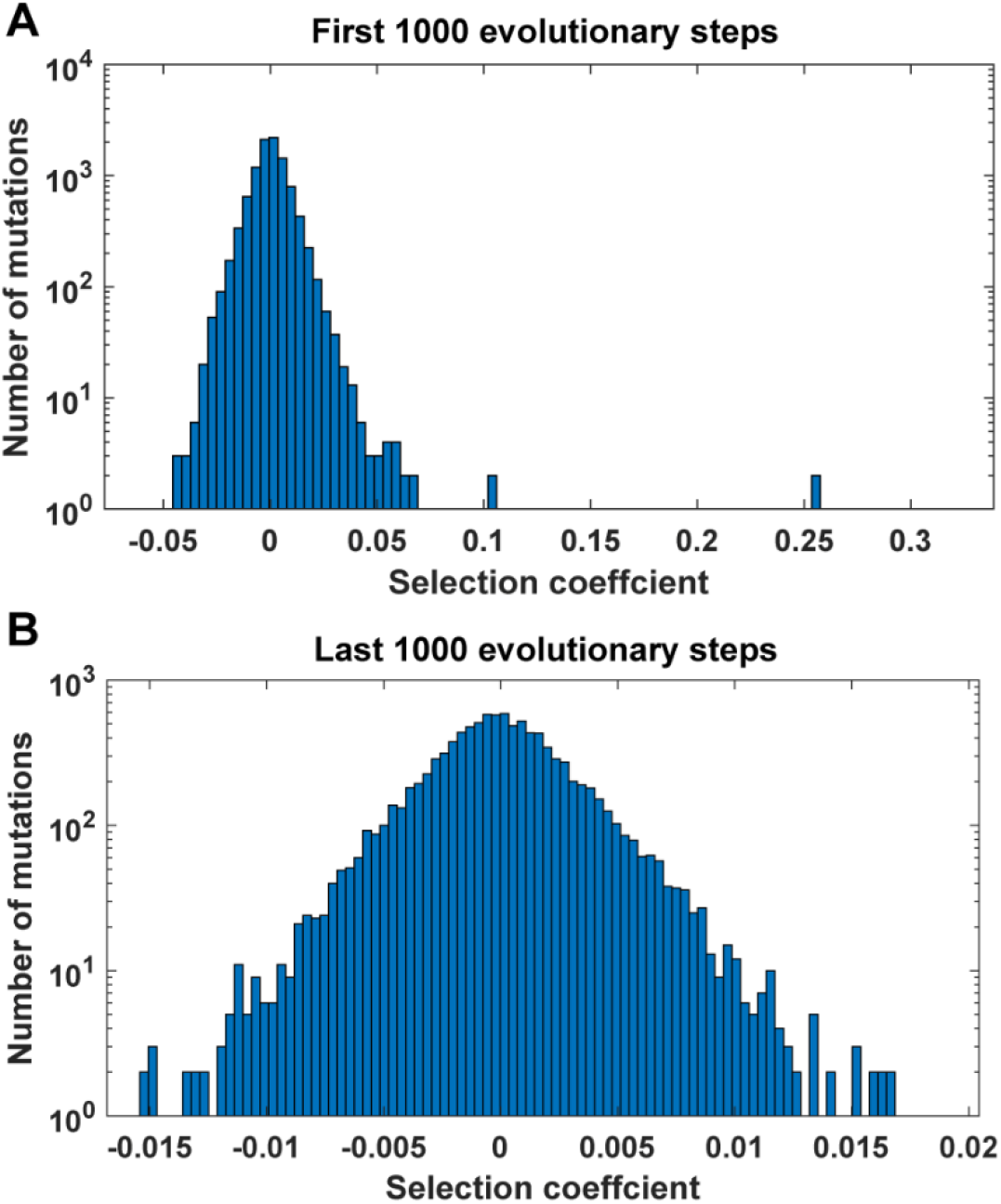
Evolutionary paths include slightly deleterious mutations. We pooled all accepted mutations from 10 evolutionary simulations under selection for high peak effector expression. Selection coefficients were calculated from the average fitness across 1,000 simulations of gene expression. Note while fitness is therefore biased by the 200 replicates used to decide to accept that mutation, this bias applies to both resident and mutant. We measure noise on top of the true distribution of fitness effects, suggesting that the underlying distribution is narrower than shown here. **(A)** Data restricted to the first 1,000 evolutionary steps, during which fitness generally increases rapidly. (**B**) Data restricted to the last 1,000 evolutionary steps, during which almost all simulations have reached a fitness plateau.

**Figure S2.**
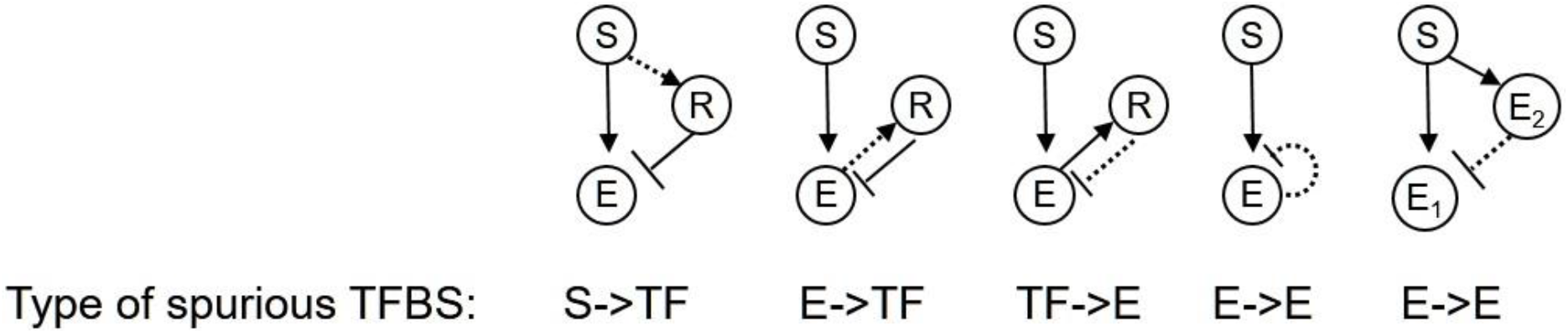
Five scenarios in which apparent but non-functional network motifs can arise from spurious TFBSs. A TFBS containing 2 mismatches can easily appear by chance in a cis-regulatory sequence, but may be deemed spurious if it has negligible functional effect. Spurious E->E TFBSs where both “Es” represent the same effector gene give rise to apparent ARs, whereas if they represent different effector proteins, they give rise to I1FFLs.

**Figure S3.**
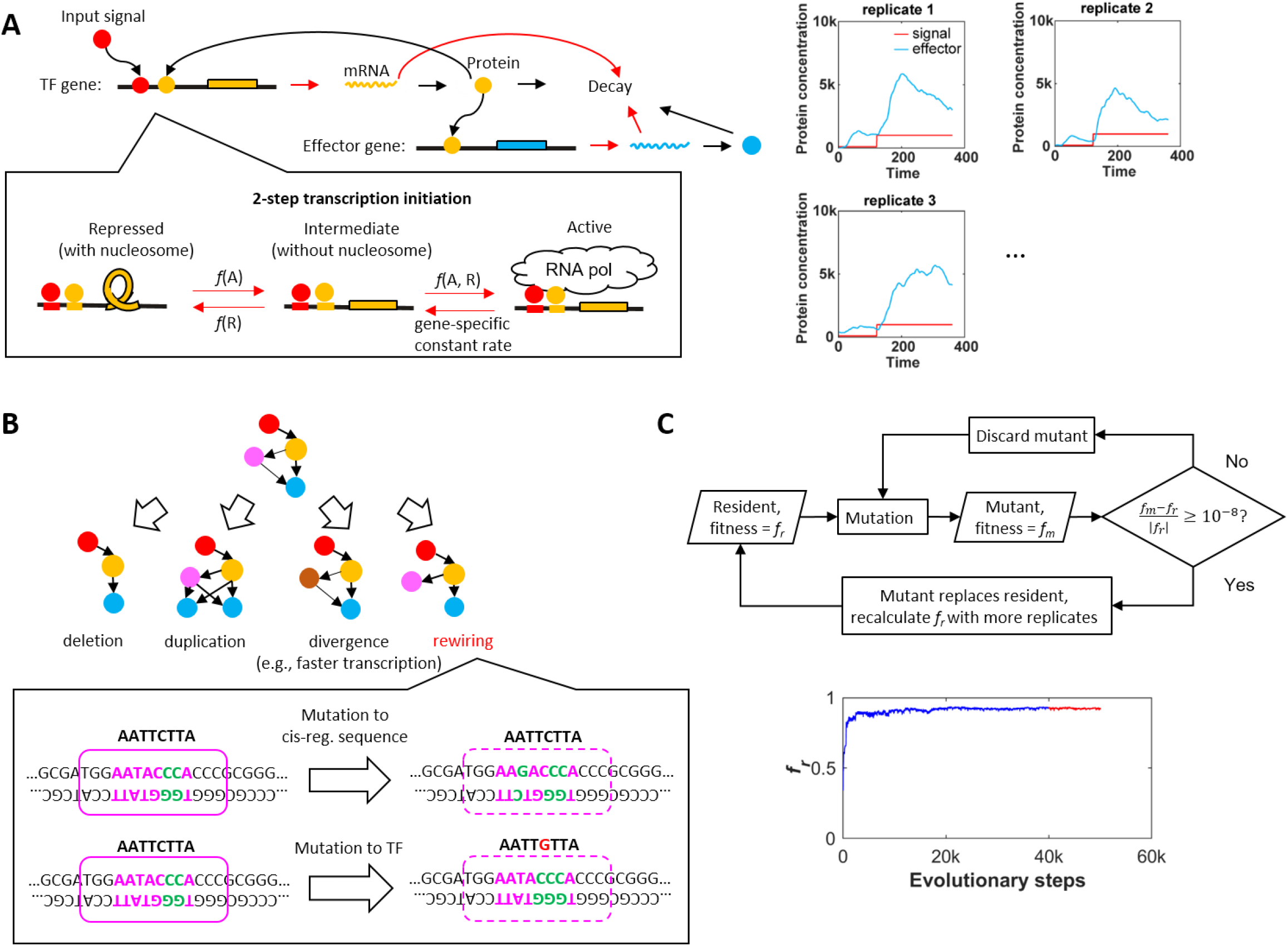
Summary of the model. **(A)** Simulation of gene expression in a TRN that has two TF genes, one of which is the effector (cyan). Here the input signal, which is simulated as an activator, binds to the cis-regulatory sequence of the non-effector TF gene (TF binding is demonstrated in **(B)**) and induces gene expression. Transcription initiation is a two-step process where most of the transition rates are functions of the concentrations of activators and/or repressors (see Transcriptional regulation in the supplement). Biological processes marked by red arrow are simulated as stochastic processes, and those marked by black arrows are simulated by solving ordinary differential equations (see Simulation of gene expression in the supplement). We use the expression levels of the effector in response to a two-stage input signal to calculate the fitness (see Methods for details). The simulation of gene expression is repeated and the average fitness of the replicates is used as the fitness of the TRN (see Methods for details). The diagram of transcription and translation is revised from Xiong et al. (2019) under a Creative Commons Attribution 4.0 International License (https://creativecommons.org/licenses/by/4.0/). **(B)** A TRN goes through one of many types of mutation (see Model Overview for details) that change the size of the network, rewire the network, or change one property of a gene in the network. The zoom-in depicts turnover of TF binding sites, which can rewire the network. The purple box represents the TF and on top of the box is the consensus binding sequence of the TF. At most two mismatches (green letters) to the consensus binding sequences can be tolerated. Point mutations in the cis-regulatory sequence of the target gene and in the consensus binding sequence of the TF can increase mismatch, causing the loss of a TF binding site. Note that the TF occupies additional sequences when binds to the DNA. **(C)** Evolution of TRNs is simulated as an origin-fixation process. Evolution starts with a random TRN, which is called the resident. if the mutant’s fitness is sufficiently high (see Methods for details), it replaces the resident and becomes the new resident (see Methods for details), which is defined as one evolutionary step. Otherwise, new mutants are generated until the replacement happens. The evolution is simulated for 50,000 evolutionary steps, which is generally long enough for the resident’s fitness to reach a plateau.

**Figure S4.**
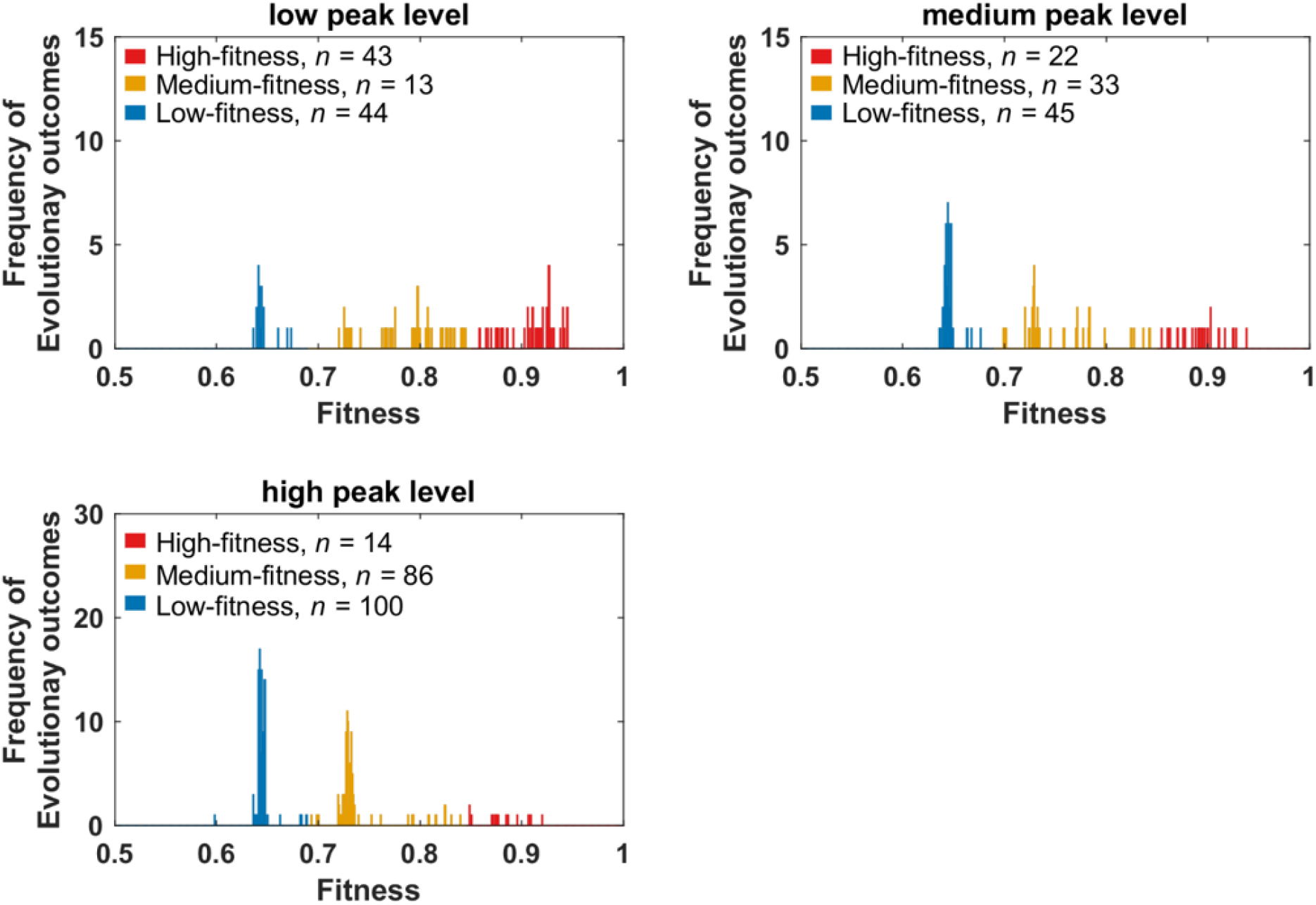
Fitness distributions of genotypes evolved with different optimal peak levels of the effector. We ran 100 evolutionary simulations for the low-peak and the medium-peak conditions, and 200 for the high-peak condition. For each simulation, we calculate the fitness of the evolved genotype as the average fitness of the last 10,000 evolutionary steps. For all three selection conditions, genotypes with fitness above 0.845 are considered as high-fitness genotypes and are further analyzed in **Fig. 2**. We used a fitness cutoff of 0.69 to separate medium-fitness genotypes and low-fitness genotypes.

**Figure S5.**
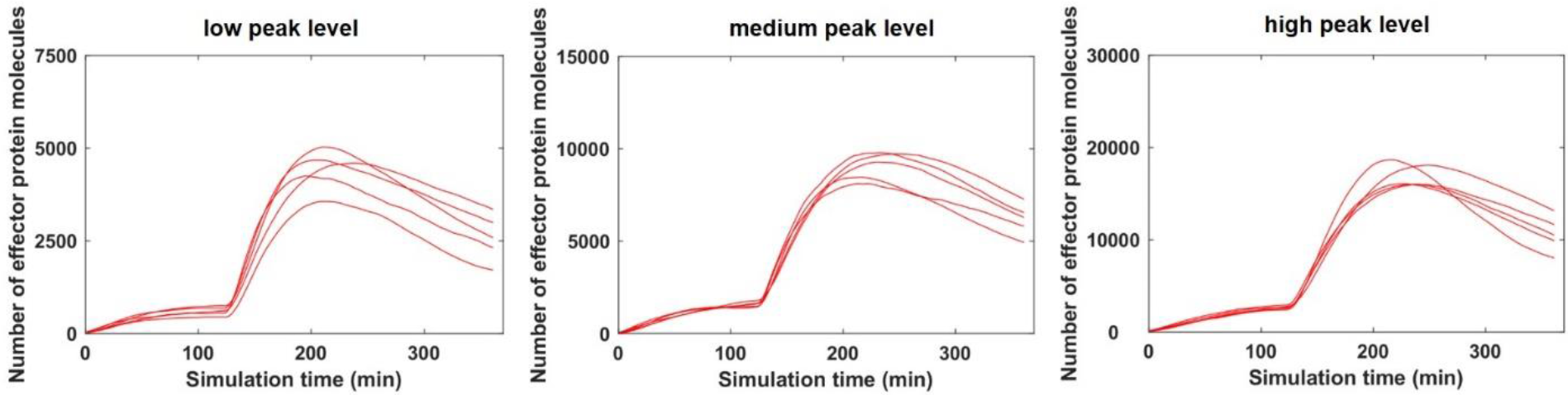
Phenotype of high-fitness replicates. For each selection condition, we randomly picked 5 high-fitness replicates from those defined in Fig. S4. We ran 200 simulations to characterize the expression profile of the effector, as found at evolutionary step 50,000 in each replicate. Each trajectory shows the expression levels of the effector averaged across the 200 simulations, and starts after the burn-in of gene expression (see **Methods** for details).

**Figure S6.**
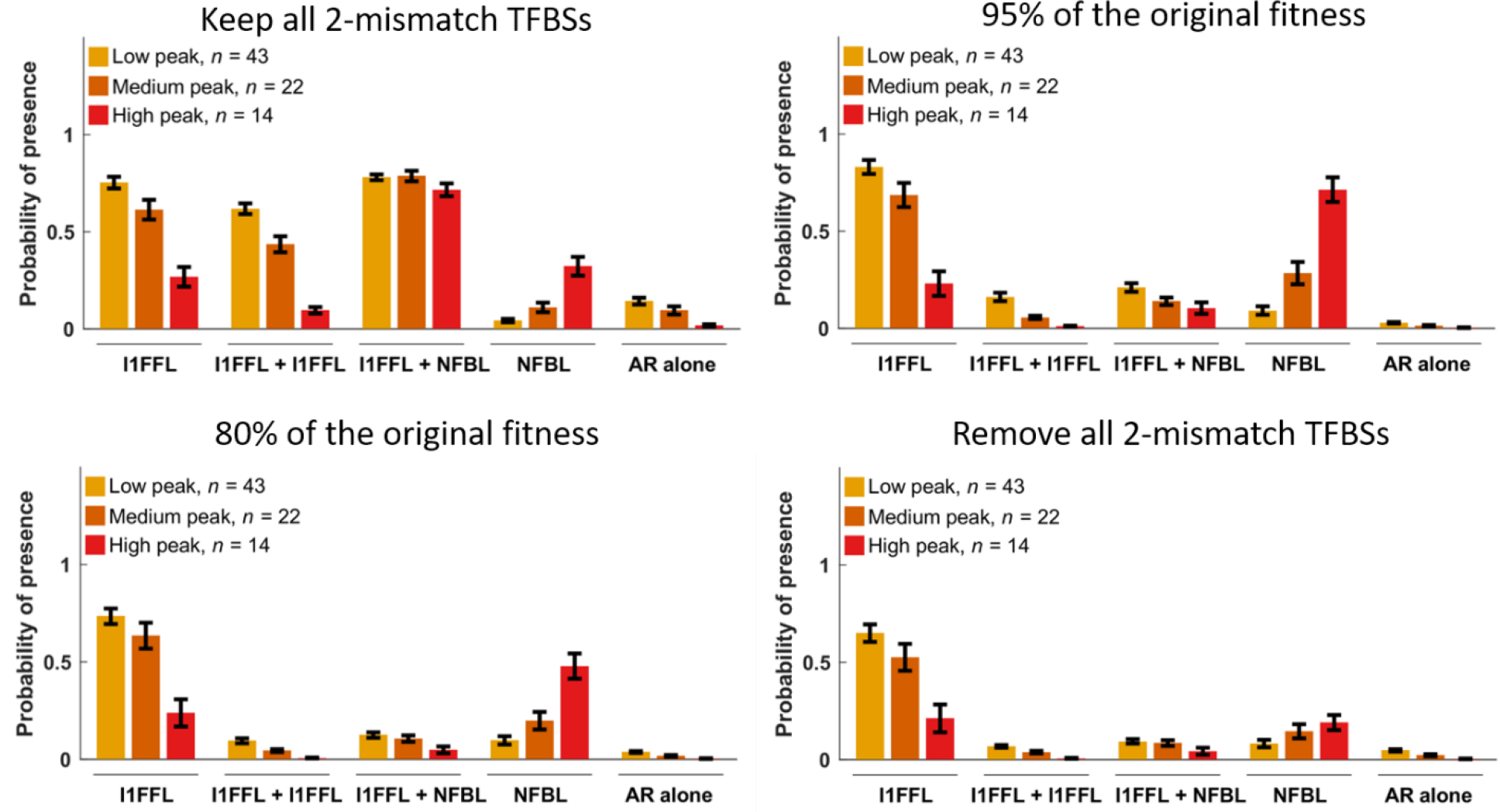
The relative occurrences of motifs do not depend strongly on the criteria for removing spurious 2-mismatch TFBSs. Results are from the same high-fitness evolutionary replicates shown in Fig. 2A, where sets of TFBSs were excluded when their removal yielded fitness of at least 99% of the fitness observed in their presence. Data are shown as mean ± SE over replicates.

**Figure S7.**
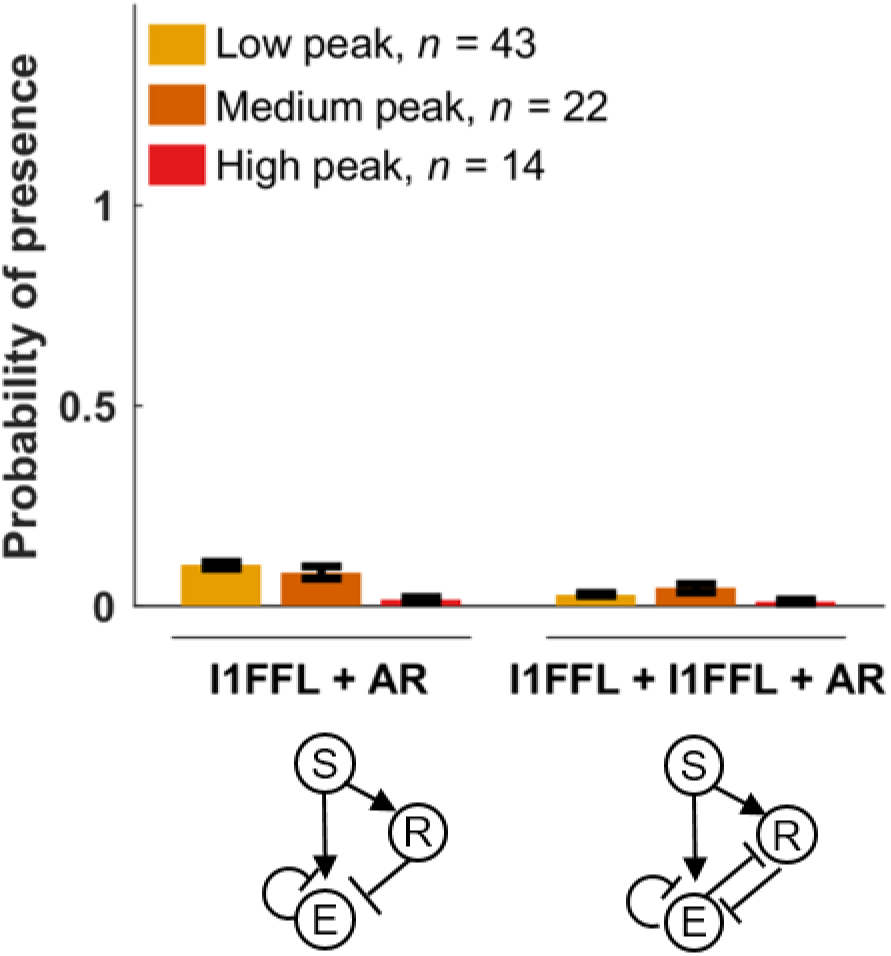
Auto-repression (AR) rarely evolves with I1FFLs or overlapping I1FFLs. In high-fitness genotypes evolved under selection for pulse generation, there were few auto-repressing effectors co-occurring with other motifs, and for simplicity, they were therefore grouped in **Fig. 2** with the motif with which they co-occurred. We note that when the repressor of an I1FFL-NFBL conjugate is an effector, this effector can form auto-repression. We classified such case as a stand-alone AR, because from the perspective of this effector, it is not regulated by an I1FFL, NFBL, overlapping I1FFL, or I1FFL-NFBL conjugate. Data are shown as mean ± SE over replicates.

**Figure S8.**
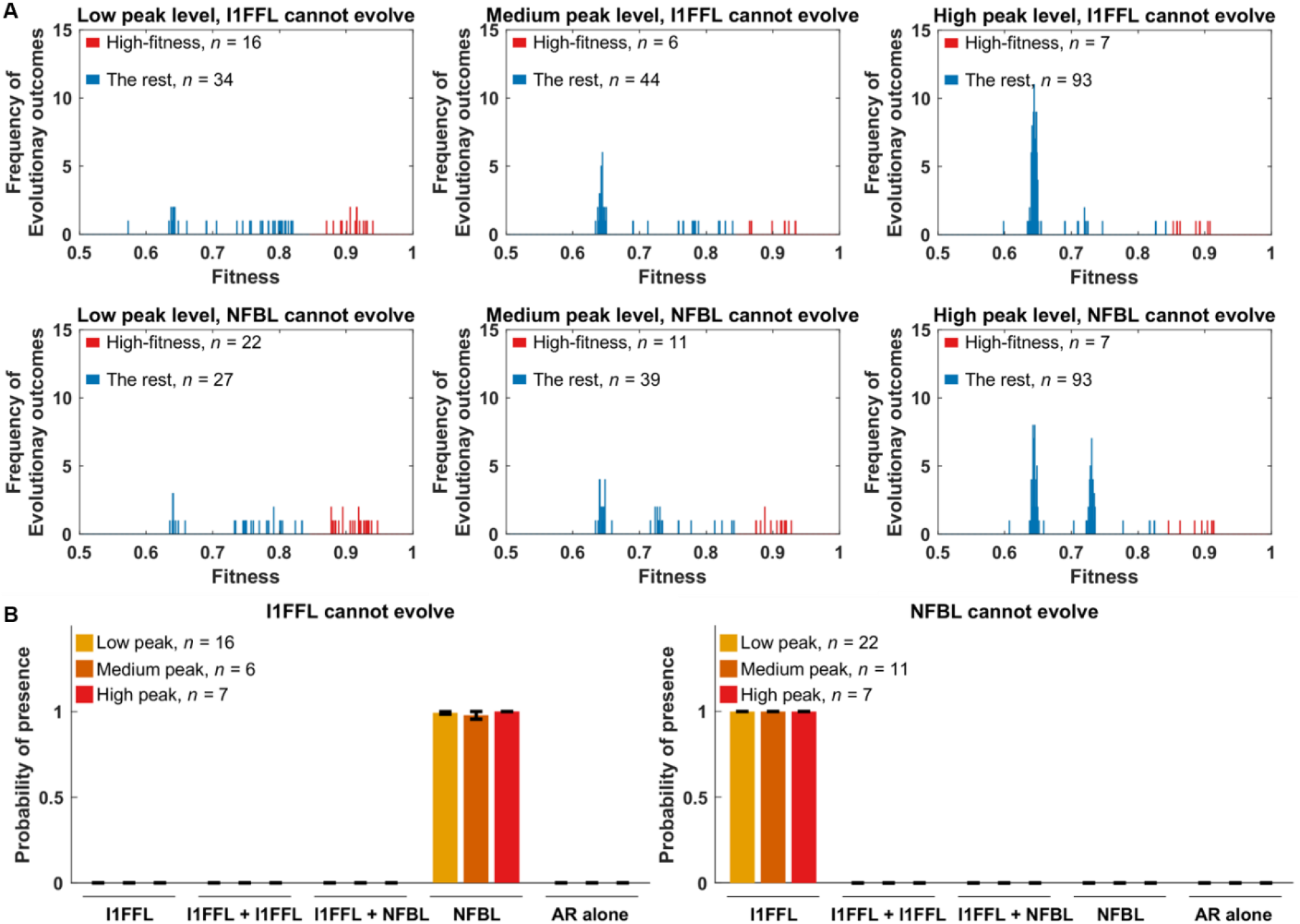
Fitness distributions and network occurrences of genotypes with restricted solutions under different selection conditions. **(A)** Each panel under low peak and medium peak selection summarized 50 evolutionary simulations, and the two panels under high peak each summarize 100 evolution simulations. Under the condition where we select for a low peak and prevent NFBLs from evolving, we removed one simulation that was terminated prematurely before evolving 50,000 evolutionary steps. This particular simulation failed to find a mutant that has higher fitness than the resident phenotype even after 2,000 trials. To classify a genotype as high-fitness (red), we apply the same fitness cutoff as in **Fig. S1**. The average fitness of the high-fitness genotypes is shown in **Fig. 3B**. See legend of **Fig. 2B** for description of modifications to prevent the evolution of NFBLs or I1FFLs. **(B)** In the high-fitness genotypes, when either I1FFL or NFBL is not allowed to evolve, the other motif almost always evolves. Data are shown as mean ± SE over replicates.

**Figure S9.**
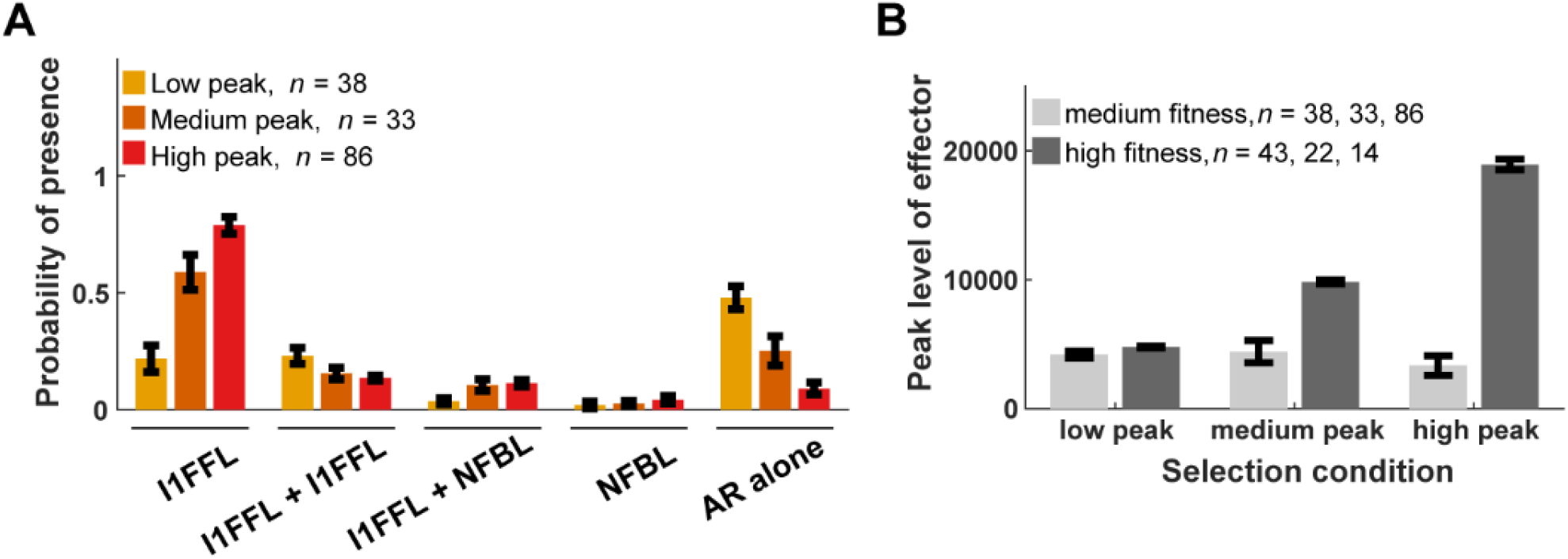
Medium-fitness genotypes fail to achieve high peak effector expression, and primarily evolve I1FFLs and auto-repression. **(A)** Methods are the same as for **Fig. 2A**, applied here to medium-fitness evolutionary replicates. **(B)** For each high-fitness and medium-fitness replicate shown in Fig. S4, we average the peak protein levels of the effector over 200 replicate simulations of gene expression. Data are shown as mean ± SE over replicates.

**Figure S10.**
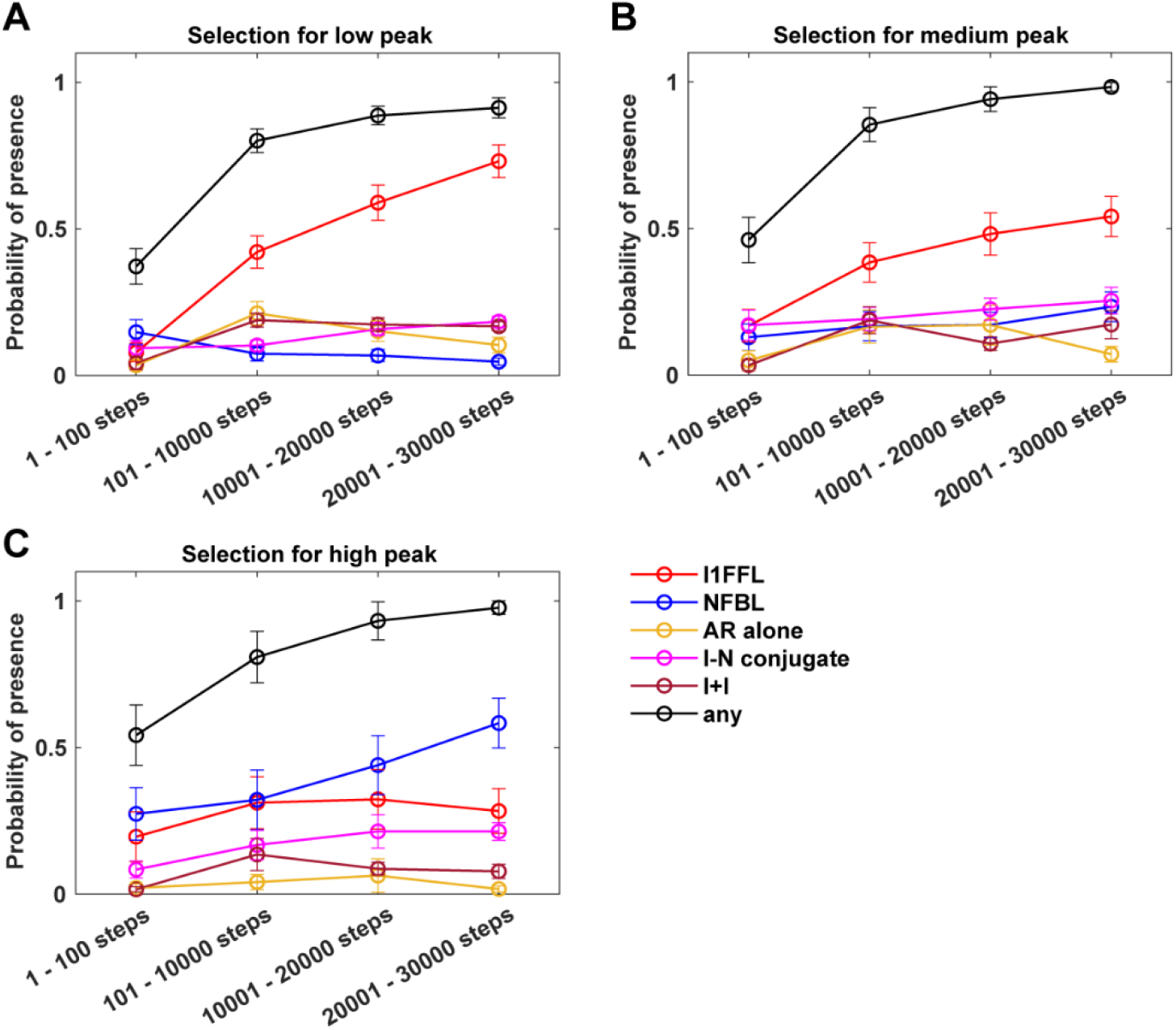
The occurrence of all motifs during evolution. We calculated the proportion of evolutionary steps that contain at least one motif of that type. Details are the same as for **Fig. 3**, except here we show a broader range of motifs as shown in **Fig. 2**. Data are shown as mean ± SE over the high-fitness replicates.

**Figure S11.**
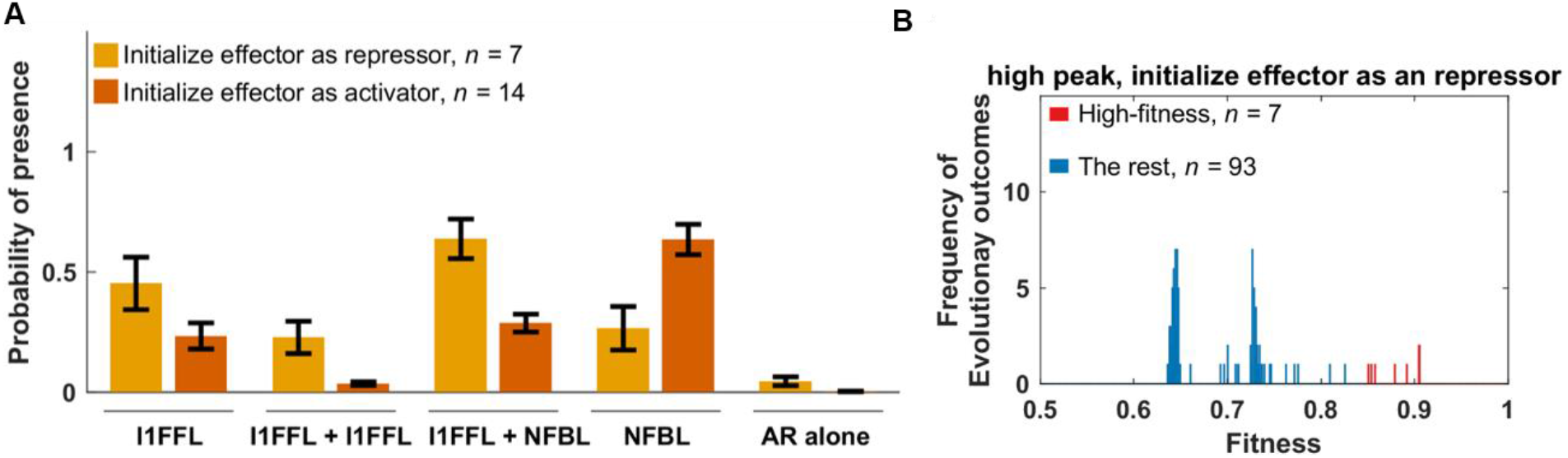
Initializing the effector as a repressor facilitates the evolution of I1FFLs. We repeated evolution under selection for high peak effector expression, but initialized the effector as a repressor rather than as an activator. **(A)** Motif occurrence compared to the activator-initialized evolutionary conditions given in **Fig. 2**. Data are shown as mean ± SE over replicates. **(B)** Fitness of the evolved TRNs. Similar to Fig. S1, TRNs with fitness of 0.845 or higher are considered high-fitness.

**Figure S12.**
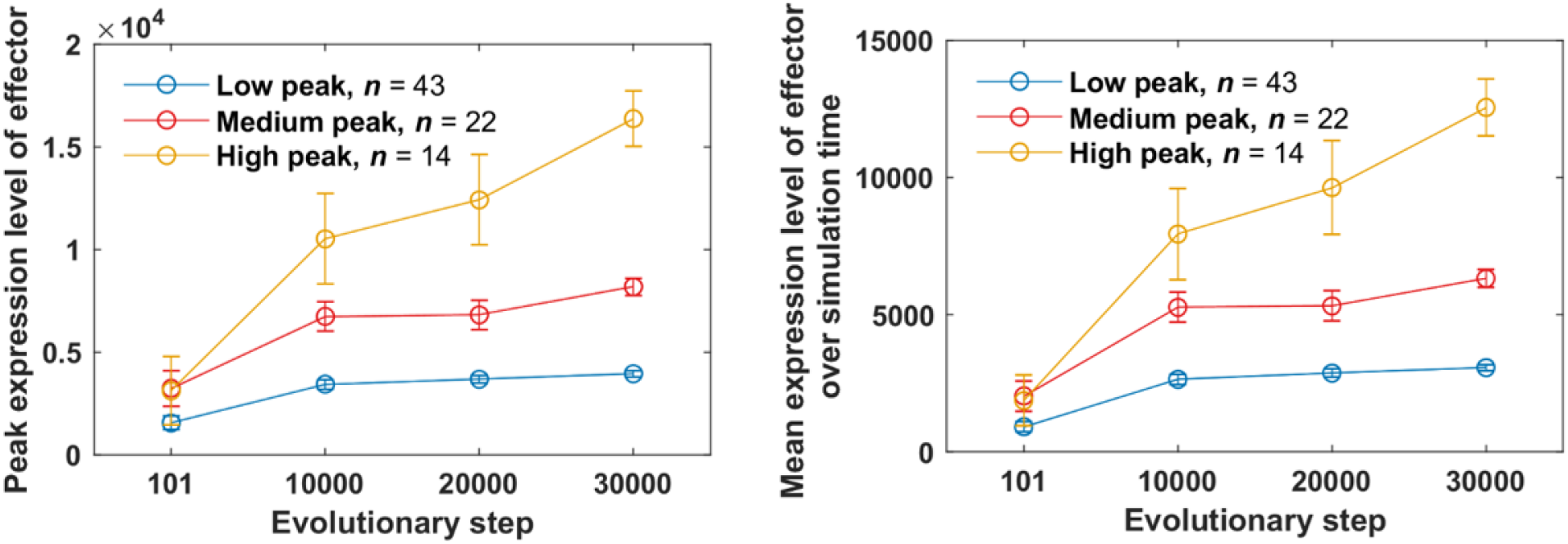
High peak effector expression evolves slowly. For each high-fitness replicate shown in Fig. 2A, we average the peak protein levels of the effector over 200 replicate simulations of gene expression. Data are shown as mean ± SE over replicates.

**Figure S13.**
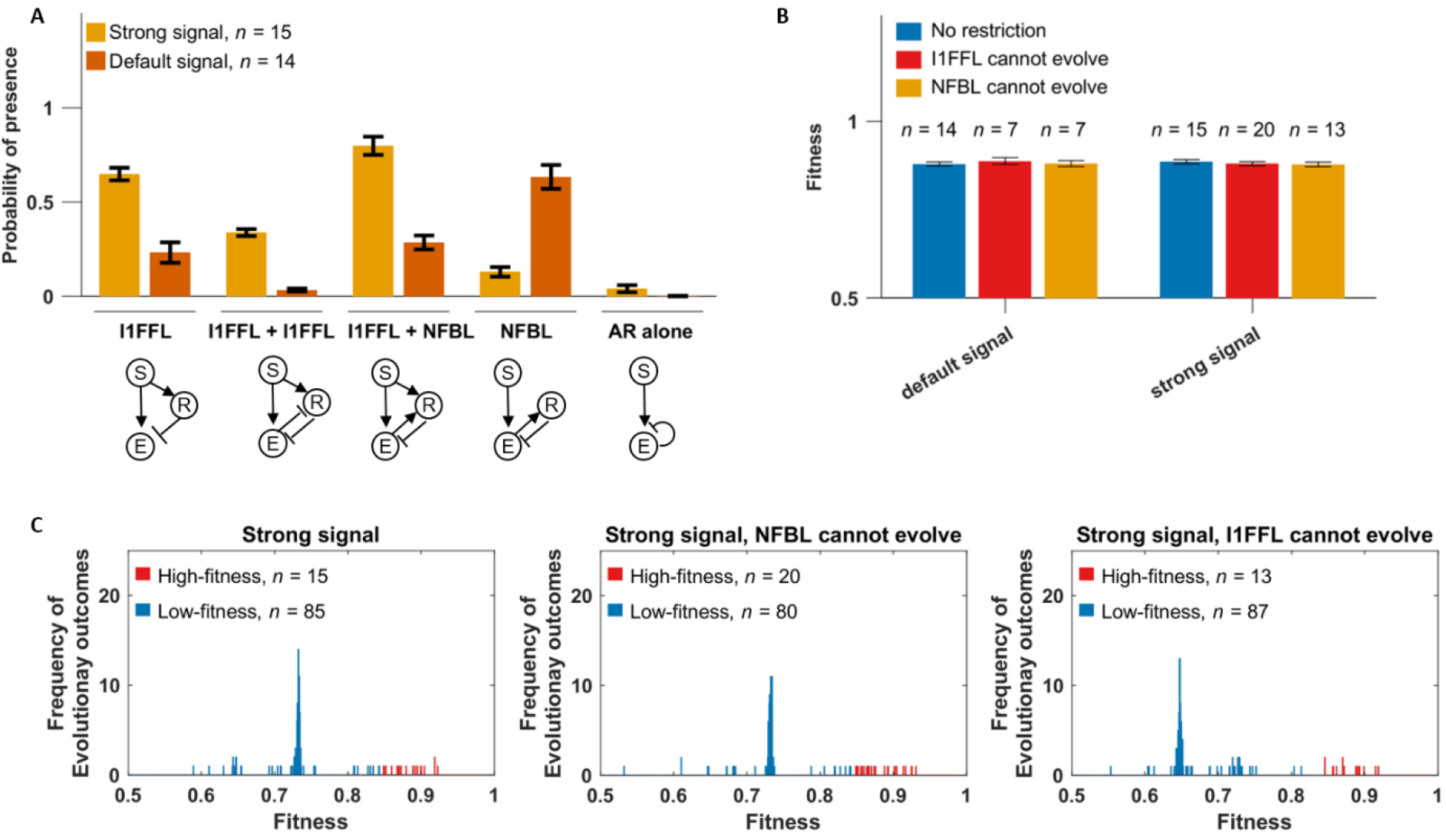
A stronger signal increases I1FFL prevalence. We compared evolution under the default signal, where signal strength increases from 100 molecules per cell to 1,000 molecules per cell, to a stronger signal, where the signal strength increases from 1,000 molecules per cell to 10,000 molecules per cell. **(A)** Occurrence of different motifs in high-fitness genotypes. **(B)** I1FFLs or NFBLs can yield similar fitness under a given signal regime. Data are shown as mean ± SE over replicates. **(C)** Fitness distribution of genotypes evolved with a strong signal without (leftmost) or with (middle and rightmost) restrictions on evolution. Similarly to Figs. S1 and S3, we define high-fitness genotypes to be those with fitness greater or equal to 0.845.

**Figure S14.**
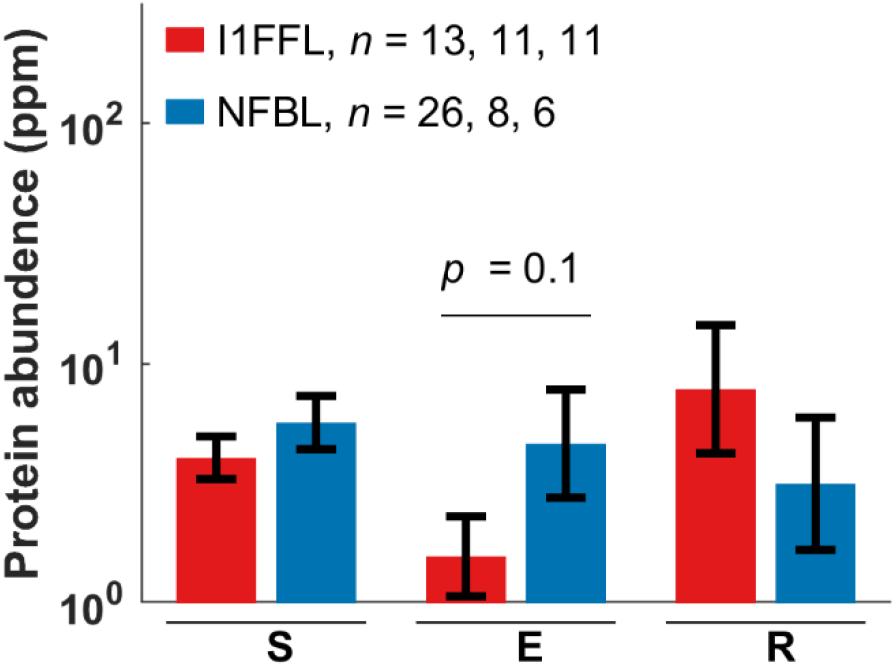
Effectors have higher protein expression in NFBLs than in I1FFLs in *S. cerevisiae* across a more comprehensive dataset. Protein expression levels are from the “GPM, Aug, 2014” dataset provided by PaxDB (Wang et al. 2015). Data are shown as mean ± SE (of log-transformed data in the case of protein expression) over each network position across all instances of the motif, excluding positions where the data are not available. For each motif type, we list the numbers of genes with available expression level data at signal nodes, effector nodes, and repressor nodes. Statistical significance is assessed using two-tailed t-tests.

## Details of the model

The following sections are copied from Xiong et al. (2019). Parts of the original text were rewritten or deleted for brevity. The original article was licensed under the Creative Commons Attribution 4.0 International License, which grants free copy and modification. A copy of the license can be found at https://creativecommons.org/licenses/by/4.0/.

### TF binding

In our model, each gene is controlled by a 150-bp cis-regulatory region, corresponding to a typical yeast nucleosome-free region within a promoter (Yuan et al. 2005). TFBSs can evolve in the cis-regulatory region, and we set the length of a consensus binding sequence to be 8 bp. Assuming that only one of the four nucleotides is a good match at each of the 8 base pairs, then the 8-bp consensus binding sequence has an information of 16 bits, which is slightly larger than that of a typical yeast TF (13.8 bits) (Wunderlich and Mirny 2009). We assume a higher information content than seen empirically in order to reduces the number of TFBSs within the cis-regulatory regions to a point that our computational power can handle. We allow up to 2 mismatches in the consensus binding sites, based on the finding that, with up to 2 mismatches in the 6-bp binding sequence, some yeast TFs can still bind DNA at above background level (Maerkl and Quake 2007). To capture competitive binding between TFs, we assume that two TFs cannot simultaneously occupy overlapping stretches, which we assume extend beyond the recognition sequence to occupy a total of 14 bp (Zhu and Zhang 1999).

We denote the dissociation constant of a TFBS with *m* mismatches as *K*_d_(*m*). Sites with *m* > 3 mismatches are assumed to still bind at a background rate equal to *m* = 3 mismatches, with dissociation constant *K*_d_(3) = 10^−5^ mole per liter (Maerkl and Quake 2007) for all TFs. We assume that each of the last three base pairs makes an equal and independent additive contribution Δ*G*_bp_ < 0 to the binding energy (Benos et al. 2002). We ignore cooperativity in binding. Dissociation constants of eukaryotic TFs for perfect TFBSs can range from 10^−5^ mole per liter (Park et al. 2004) to 10^−11^ mole per liter (Nalefski et al. 2006). We initialize each TF with its own value of log_10_(*K*_d_(0)) sampled from a uniform distribution between −6 and −9, with mutation capable of further expanding this range, subject to *K*_d_(0) < 10 ^5^ mole per liter. Substituting *m* = 0 and *m* = 3 into

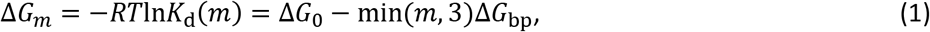

where *R* is the gas constant and *T* is temperature, we can solve for *ΔG*_bp_ and *ΔG*_0_, and thus obtain *K*_d_(1) and *K*_d_(2) (the dissociation constants for TFBS with one and two mismatches, respectively).

We rescale *K*_d_ values to effective *K*_d_ values to account for the “dilution” of TFs by non-specific TF binding sites (NSBSs) in the genome. A haploid *S. cerevisiae* genome is 12 Mb, 80% of which is wrapped in nucleosomes (Lee et al. 2007), yielding approximately 10^6^ potential NSBSs. In a yeast nucleus of volume 3 × 10^−15^ liters, the NSBS concentration is of order 10^−4^ mole per liter. To find the concentration of free TF [TF] in the nucleus given a total nucleic TF concentration of *C_TF_*, we consider

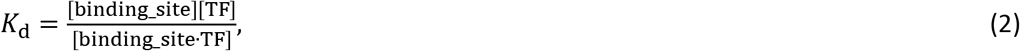

in the context of NSBSs, substitute [TF∙NSBS] with *C*_TF_ – [TF], and solve for

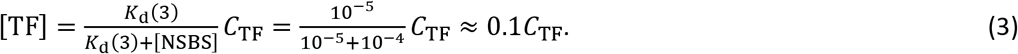

Thus, about 90% of total TFs are bound non-specifically, leaving about 10% free. The relatively small number of specific TFBSs is not enough to significantly perturb the proportion of free TFs, and so for the specific TFBSs with *m* < 3 that are of interest in our model, we simply use 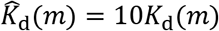 to account for the reduction in the amount of available TF due to non-specific binding. We also convert 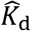 from the units of mole per liter in which *K*_d_ is estimated empirically to the more convenient molecules per nucleus. The rescaling factor *r* for which 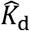 (in molecule per nucleus) 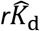 (in mole per liter) is 3 ×10^−15^ liter per nucleus × 6.02 × 10^23^ molecule mole^−1^ = 1.8 × 10^9^ molecule cell^−1^ liter mole^−1^. Taken together, 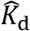 (molecule per nucleus) = 10*rK*_d_ (mole per liter), where the factor 10 accounts for non-specific TF binding.

### TF occupancy

Here we calculate the probability that there are *A* activators and *R* repressors bound to a given cis-regulatory region at a given moment in gene expression time. First we note that if we consider TF *i* binding to TFBS *j* in isolation from all other TFs and TFBSs, Supplementary Equation 4 gives us the probability of being bound:

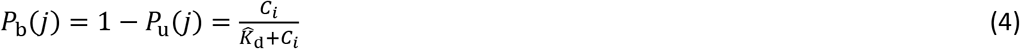

Let 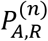 be a term proportional (for a given value of *n*) to the combined probability of all binding configurations in which exactly *A* activators and *R* repressors are bound to the first *n* binding sites along the cis-regulatory sequence. We calculate 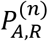 recursively, considering one additional TFBS at each step. Note that if two different TFs bind to exactly the same location on a cis-regulatory region, we treat this as two TFBSs, not as one, and treat first one and then the other in our recursive algorithm.

Consider the case where the (*n*+1)^th^ binding site belongs to an activator. The case where this activator is not bound contributes 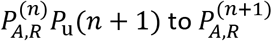. If it is bound, then we must also take into account that the (*n*+1)^th^ binding site overlaps (partially or completely) with the last *H* ≥ 0 sites, and so contributes 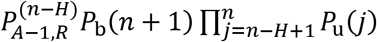. Taken together,

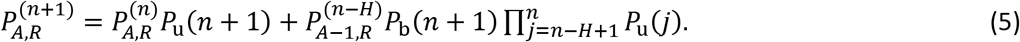

Similarly, if the (*n*+1)^th^ site belongs to a repressor, we have

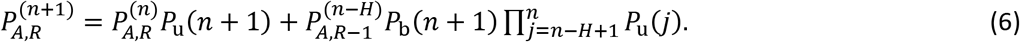

By definition, 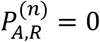 for binding configurations that are impossible, e.g. those with negative *A* or negative *R*. We initialize the recursion at *n* = 0, where the only valid binding configuration is for *A* = *R* = 0, i.e. 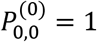. At *n* = 1, 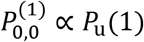 and if the binding site belongs to an activator 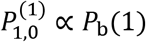; otherwise, 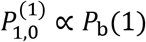. For a gene where the total number *N* of TFBSs is 1, 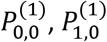, and 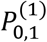 sum to 1 and normalization is unnecessary. For higher values of *N* = *N*_Act_ + *N*_Rep_ TFBSs, where *N*_Act_ and *N*_Rep_ are the total numbers of activator binding sites and repressor binding sites, respectively, we normalize 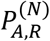 at the end of the recursion by dividing by 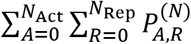 to get the probability of binding configurations that include exactly *A* activators and *R* repressors.

#### *r*_Act_to_Int_

Transcription initiation over an interval of time *r*_transc_init_ is proportional to the proportion of time spent in the Active state. Assuming a steady state between <u>Rep</u>ressed, <u>Int</u>ermediate, and <u>Act</u>ive states, as a function of current TF concentrations, we have:

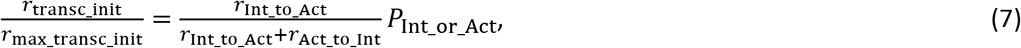

where *P*_Int_or_Act_ is the probability a gene is at Intermediate or Active. We set *r*_max_transc_init_ (the rate of transcription given 100% Active state) to 6.75 min^−1^, based on the corresponding rate when a model of the *PHO5* promoter is fit to data (Brown et al. 2013). In this model fit, the constitutively expressed *PHO5* promoter is free of nucleosomes 80% of the time, i.e. *P*_Int_or_Act_ = 0.8. We take these two values as universal for constitutively expressed genes, and assume that variation in *r*_Act_to_Int_ is responsible for variation in *r*_transc_init_. To identify a set of constitutively expressed genes, we identified 225 genes that have mRNA production rate of at least 0.5 molecule min^−1^ from genome-wide measurements (Pelechano et al. 2010); this threshold corresponds to low H2A.Z occupancy (Guillemette et al. 2005). We set *r*_transc_init_ to the production rate of mRNA of these 225 genes, and solve for gene-specific *r*_Act_to_Int_ from Eq. S7. We fit the solutions to a log-normal distribution and arrive at 10^N(1.27,0.226)^ min^−1^.

To initialize values of *r*_Act_to_Int_ for each gene, we sample from this distribution. We also set lower and upper bounds for allowable values; if either the initial sample or subsequent mutation put *r*_Act_to_Int_ beyond these bounds, we set the value of *r*_Act_to_Int_ to equal to boundary value. We set the lower bound for *r*_Act_to_Int_ at 0.59 min^−1^, half the minimum of the values inferred from the set of 225 genes. To set an upper bound, we use the low H2A.Z occupancy bound of *r*_transc_init_ = 0.5, which gives a solution of 32.34 min^−1^; we double this to set the upper bound as 64.7 min^−1^.

### Transcription delay times

Yeast protein lengths fit a log-normal distribution of 10^N(2.568, 0.34)^ amino acids (from the Saccharomyces Genome Database (SGD Project), excluding mitochondrial proteins; YeastMine (Balakrishnan et al. 2012) was used to query the database and to download data). We sample ORF length *L* from this distribution. To constrain the values of *L*, we set a lower bound of 50 amino acids and an upper bound of 5,000 amino acids; the longest protein in SGD is 4910 amino acids. If either initialization or mutation put *L* beyond these bounds, we set the value of *L* to the boundary value.

With an mRNA elongation rate of 600 codon per min (Larson et al. 2011; Hocine et al. 2013), it takes *L* / 600 minutes to transcribe the ORF of an mRNA. Also including time for transcribing UTRs and for transcription termination, and ignoring introns for simplicity, it takes 290 seconds to complete transcription of the yeast *GLT1* gene (Larson et al. 2011), whose ORF is 6.4kb. Putting the two together, we infer that transcribing the UTRs and terminating transcription takes around 1 minute for *GLT1*.

Generalizing to assume that transcribing UTRs and terminating transcription takes exactly 1 minute for all genes, producing an mRNA from a gene of length *L* takes 1 + *L* / 600 minutes.

### Translation delay times and *r*_protein_syn_

We model a second delay between the completion of a transcript and the production of the first protein from it. The delay comes from a combination of translation initiation and elongation; it ends when the mRNA is fully loaded with ribosomes all the way through to the stop codon and the first protein is produced. We ignore the time required for mRNA splicing; introns are rare in yeast (Dujon 1996). mRNA transportation from nucleus to cytosol, which is likely diffusion-limited (Niño et al. 2013; Smith et al. 2015), is fast even in mammalian cells (Mor et al. 2010) let alone much smaller yeast cells, and the time it takes is also ignored. The median time in yeast for initiating translation is 0.5 minute (Table 1 in Siwiak et al. 2010), and the genomic average peptide elongation rate is 330 codon/min (Siwiak et al. 2010). After an mRNA is produced, we therefore wait for 0.5 + *L* / 330 minutes, and then model protein production as continuous at a gene-specific rate *r*_protein_syn_.

To calculate *r*_protein_syn_, we combine the gene-specific ribosome densities *D* along the mRNAs and the gene-specific peptide elongation rates *E*, both measured in yeast (Siwiak et al. 2010). The values of *DE* across yeast genes fit the log-normal distribution 10^N(0.322, 0.416)^ molecule mRNA^−1^ min^−1^; we initialize *r*_protein_syn_ for each gene by sampling from this distribution. We set the lower bound for *r*_protein_syn_ at half the minimum observed value of *DE* (4.5 × 10^−3^ molecule mRNA^−1^ min^−1^). The upper bound corresponds to an mRNA fully occupied by rapidly moving ribosomes. Each ribosome occupies about 10 codons (Siwiak et al. 2010), and the peptide elongation rate can be as high as 614 codon per min (Waldron et al. 1977). If ribosomes are packed closely together at 10 codons apart, a protein comes off the end of production in the time taken to elongate 10 codons, i.e. proteins are produced at 61.4 molecules per minute. If either initialization or mutation put *r*_protein_syn_ beyond these bounds, we set the value of *r*_protein_syn_ to the boundary value.

### mRNA and protein decay rates

We fit a log-normal distribution 10^N(−1.49, 0.267)^ min^−1^ to yeast mRNA degradation rates (Wang et al. 2002), and initialize the mRNA degradation rate *r*_mRNA_deg_ for each gene by sampling from this distribution. We set lower and upper bounds for *r*_mRNA_deg_ at half the minimum and twice the maximum observed values (7.5 × 10^−4^ min^−1^ and 0.54 min^−1^), respectively. If either initialization or mutation put *r*_mRNA_deg_ beyond these bounds, we set the value of *r*_mRNA_deg_ to the boundary value.

Expressing the estimated half-lives of yeast proteins (Belle et al. 2006) in terms of protein degradation rates, they fit the log-normal distribution 10^N(−1.88, 0.56)^ min^−1^; we initialize gene-specific protein degradation rates *r*_protein_deg_ by sampling from this distribution. We ignore the additional reduction in protein concentration due to dilution as the cell grows and thus increases in volume. We set lower and upper bounds for *r*_protein_deg_ at half the minimum and twice the maximum observed degradation rate (3 × 10^−6^ min^−1^ and 0.69 min^−1^), respectively. If either initialization or mutation put *r*_protein_deg_ beyond these bounds, we set the value of *r*_protein_deg_ to the boundary value.

### Simulation of gene expression

Our algorithm is part-stochastic, part-deterministic. We use a Gillespie algorithm (Gillespie 1977) to simulate stochastic transitions between <u>Rep</u>ressed, <u>Int</u>ermediate, and <u>Act</u>ive chromatin states, and to simulate transcription initiation and mRNA decay events. We refer to these as “Gillespie events”. The completion of transcription to produce a complete mRNA, and subsequent ribosomal loading onto the mRNA, are referred to as “fixed events” (they require fixed times of 1 *+ L* / 600 minutes and 0.5 + *L* / 330 minutes, respectively). Scheduled changes in the strength of the external signal are also fixed events. Protein production and degradation are described deterministically with ODEs, and updated frequently in order to recalculate TF concentrations and hence chromatic transition rates. Updates occur at the time of Gillespie and fixed events, and also in between as described later below.

The total rate of all Gillespie events is

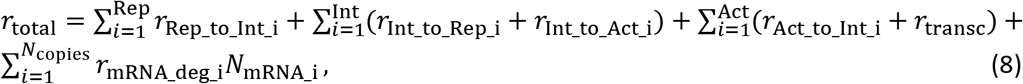

where Rep, Int, and Act are the numbers of gene copies in our haploid model that are in the Repressed, Intermediate, and Active chromatin states, respectively, *N*_mRNA_i_ is the number of completely transcribed mRNA molecules from gene *i*, and *N*_copies_ is the total number of gene copies. We only simulate degradation of full transcribed mRNA, and not that of mRNA that are still being transcribed, because the latter are already captured implicitly by *r*_max_transc_init_, which is based on mRNAs that complete transcription (Brown et al. 2013). Once an mRNA finishes transcription, it is subjected to degradation regardless of whether ribosome loading is complete.

The waiting time Δ*t*_G_ before the next Gillespie event is

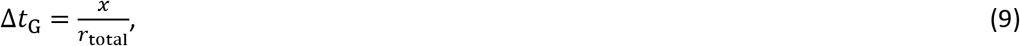

where *x* is random number drawn from an exponential distribution with mean 1. Which Gillespie event takes place next is sampled only if a different update does not happen first. If a fixed event is scheduled to happen first at *Δt*_F_ < *Δt*_G_, we advance time by *Δt*_F_, update the state of the cell, and calculate a new *r*_total_′. Since the cellular activity has been going on with the old rate *r*_total_ for *Δt*_F_, the remaining “labor” required to trigger the Gillespie event planned earlier is reduced. The new waiting time, *Δt*_G_′, to trigger the planned Gillespie event is

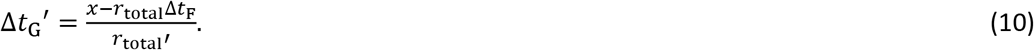

Gene duplication creates *n* ≥ 1 genes copies producing the same protein, where each copy *i* might have diverged in their production rate *r*_protein_syn_i_ and degradation rate *r*_protein_deg_i_. Complete proteins are produced continuously once an mRNA molecule is fully loaded with ribosomes, which occurs 0.5 + *L* / 330 minutes after transcription is complete – the concentration of such molecules is denoted *N*_mRNA_aft_delay_i_(*t*). The total concentration of a protein obeys:

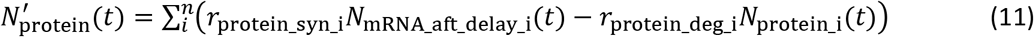

Protein concentrations are updated using a closed-form integral of Supplementary Equation 11

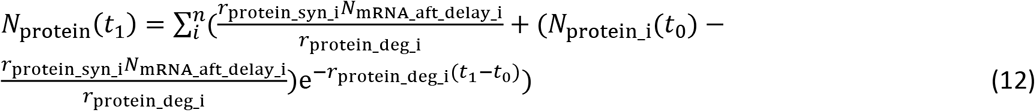

with this expression updated every time a Gillespie or fixed event at time *t_1_* changes the value of *N*_mRNA_aft_delay_i_.

In between updates, values of *P*_A_, *P*_R_, *P*_A_no_R_, and *P*_notA_no_R_, and hence chromatin transition rates, are calculated under the approximation of constant *N*_protein_. Additional updates, above and beyond fixed and Gillespie events, are performed in order to ensure that chromatin transition rates do not change too dramatically from one update to the next. We use a target of *D* = 0.01 for the amount of change tolerated in the values of *P*_A_, *P*_R_, *P*_A_no_R_, and *P*_notA_no_R_, in order to schedule updates after time Δ*t*_U_, which are triggered when neither a Gillespie event nor a fixed event occurs before this time has elapsed, i.e. when Δ*t*_U_ < Δ*t*_F_ and Δ*t*_U_ < Δ*t*_G_.

There is the greatest potential for large changes after an update that changes the value of *N*_mRNA_aft_delay_i_. In this case, we solve for the time interval for which the probability that TF *i* would be bound to a single perfect and non-overlapping TFBS would change by *D*, by choosing Δ*t*_U_ > 0 that satisfies

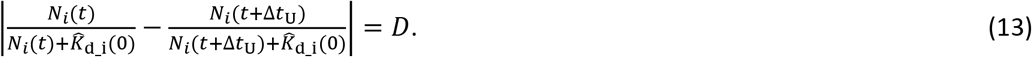

where the two left-hand terms are derived from Supplementary Equation 4. A solution for Δ*t*_U_ may not exist, e.g. if the concentration of TF *i* is decreasing but *P*_b_i_(*t*) < *D*. In such cases, we set Δ*t*_U_ to infinity.

When the previous update does not change any *N*_mRNA_aft_delay_i_ values, then we modify Δ*t*_U_ adaptively. Let *d* be the maximum of Δ*P*_A_, Δ*P*_R_, Δ*P*_A_no_R_, and Δ*P*_notA_no_R_ during the last update, and Δ*t* be the advance in time between the last two updates. We then schedule an update at

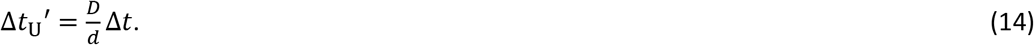

After an update that changes the value of *N*_mRNA_aft_delay_i_, we use the smaller value from Supplementary quations 13 and 14. These additional update times are discarded and recalculated when a Gillespie or fixed event occurs first. Supplementary Figure 12 of Xiong et al. (2019) shows that simulations rarely exceed the target of *D* = 0.01, and do so only modestly.

### Cost of gene expression

The cost of gene expression comes from some combination of the act of expression and from the presence of the resulting gene product. Yeast cells with plasmids carrying fast-degrading GFP had as much growth impairment as those carrying wild-type GFP (Fig. 3 of Kafri et al. 2016), suggesting that the former cost dominates. Universal costs stemming from the act of gene expression include the consumption of energy (Wagner 2005; Wagner 2007) and the opportunity cost of not using ribosomes to make other gene products (Scott et al. 2014). While some costs arise from transcription (Kafri et al. 2016), we simplify our model by attributing all of the cost of expression to the act of translation.

Kafri et al. (2016) reported that, in rich media, the growth rate of haploid yeast is reduced by about 1% when mCherry is expressed to about 2% of proteome. Setting the growth rate of the yeast when mCherry is not expressed, i.e. the fitness, to one, we have the cost of gene expression equal to 0.01. Next, we estimate the production rate of mCherry in Kafri et al. (2016) by assuming that mCherry is in steady state between production and dilution due to cell division; fluorescent proteins tend to be stable such that degradation can be ignored (Snapp 2009). Ghaemmaghami et al. (2003) estimated that a haploid yeast cell contains about 5 × 10^7^ protein molecules, 2% of which are now mCherry. Over a 90 minute cell cycle in Kafri et al. (2016), about 5 × 10^5^ mCherry molecule per cell need to be expressed in order to double in numbers. This yields a production rate of about 5 × 10^3^ mCherry molecules per minute per cell. Because the total cost of gene expression is 0.01, the cost at a protein production rate of one mCherry molecule per minute per cell, *c*_transl_, is 2 × 10^6^. Long genes should be more expensive to express than short ones; for a gene of length *L*, we assume its cost of expression is *c*_transl_*L* / 370, where 370 is the geometric mean length of a yeast protein as described above in “Transcription delay times”. Results using the length of mCherry instead, i.e. a slightly higher cost of expression of *c*_transl_*L* / 236, are unlikely to be significantly different.

The overall cost of gene expression at time *t*, *C*(*t*) is:

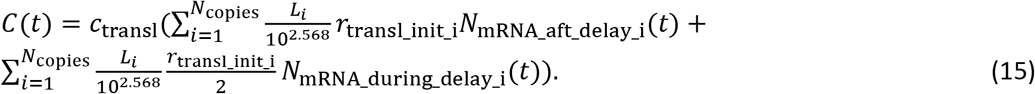

The second term represents transcripts that are on average half-loaded with ribosomes, and hence experiencing on average half the cost of translation. We integrate *C*(*t*) within segments of constant *C*(*t*) to obtain the overall cost of gene expression during a simulation.

### Mutation

Because we use an origin-fixation approach, only the relative and not the absolute values of our mutation rates matter. In *S. cerevisiae,* the rates of small indels and of single nucleotide substitutions have been estimated as 0.2 × 10^−10^ per base pair and 3.3 × 10^−10^ per base pair, respectively (Lynch et al. 2008). Thus, cis-regulatory sequences are primarily shaped by single nucleotide substitutions. We do not model small indels in the cis-regulatory sequence, but increase the single nucleotide substitution up to 3.5 × 10^−10^ per base pair to compensate. This corresponds to a rate of 5.25 × 10^−8^ per 150 bp cis-regulatory sequence.

Lynch et al. (2008) also report a rate of gene duplication of 1.5 × 10^−6^ per gene and of deletion of 1.3 × 10^−6^ per gene (not including non-deletion-based loss of function mutations). These values turned out to swamp the evolution of TFBSs and hence significantly slow down our simulations, so we chose values 10-fold lower, making both gene duplication and gene deletion occur at rate 1.5 × 10^−7^ per gene. This preserves their numerical excess but reduces its magnitude.

Our model contains 8 gene-specific parameters, namely *L*, *r*_Act_to_Int_, *r*_protein_deg_, *r*_protein_syn_, *r*_mRNA_deg_, the *K*_d_(0) of a TF, whether a TF is an activator vs. repressor, and the consensus binding sequence of a TF. We assume mutations to *L* are caused by relatively neutral small indels, which we assume to be 20% of all small indels; mutation to *L* therefore occurs at rate 1.2 × 10^−11^ per codon, i.e. 1.2 × 10^−11^*L* for a gene of length *L*. For *r*_Act_to_Int_, we assume that it is altered by 10% of all the point mutations (single nucleotide substitution and small indels) to the core promoter of a gene. The length of a core promoter is about 100 bp and is relatively constant among genes (Roy and Singer 2015), yielding a mutation rate of *r*_Act_to_Int_ of 3.5 × 10^−9^ per gene.

The remaining 6 gene-specific parameter mutation rates are parameterized with lower accuracy due to lack to data; the principal decision is which to make dependent vs. independent of gene length. TF binding to DNA depends on particular peptide motifs whose length is likely independent of TF length, therefore we make mutation rates independent of gene length for mutations to *K*_d_(0), to the consensus binding sequence of a TF, and to the activating vs repressing identity of a TF. We set the rate of each of the three mutation types to 3.5 × 10^−9^ per gene. In contrast, because the stability of an mRNA mainly depends on its codon usage (Cheng et al. 2017) and thus more codons means more opportunities for change, we assume the rate of mutation to *r*_mRNA_deg_ does depend on gene length, as do mutations to protein stability *r*_protein_deg_. *r*_protein_syn_ is determined by the density of ribosomes on mRNA and the elongation rate of ribosomes, and therefore is affected both by ribosome loading speed and by slow spots forming queues in the mRNA. Ribosome loading often relies on the 5’UTR of mRNA (Hinnebusch 2011), and 5’UTR length is positively correlated with ORF length (Tuller et al. 2009). Slow-spots in mRNA can be due to secondary structure or to suboptimal codons, therefore are also more likely to appear by mutation to long mRNAs, so we assume the rate of mutation to *r*_protein_syn_ depends on gene length. We set the mutation rates of *r*_protein_deg_, *r*_protein_syn_, and *r*_mRNA_deg_ each to 9.5 × 10^−12^ per codon; in other words, each mutation rate is 3.5 × 10^−9^ for a yeast gene of average length (on a log-scale) 10^2.568^ = 370 codons.

*r*_Act_to_Int_, *r*_protein_syn_, *K*_d_(0), *r*_protein_deg_, and *r*_mRNA_deg_ evolve as quantitative traits. They are assumed to have, in the absence of selection, a log-normal stationary distribution with mean *μ* and standard deviation *σ*, with values estimated below and listed in Supplementary Table 2. Denote the values of a parameter as *x* before mutation and *x*′ after mutation; mutation takes the form:

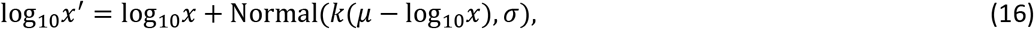

where *k* controls the speed of regressing back to the stationary distribution; we set *k* = 0.5 for all 5 parameters. To set values of *μ*, central tendency estimates of these five values (from Supplementary Table 1) are adjusted according to our expectations about mutation bias. We assume a mutation bias toward faster mRNA degradation *r*_mRNA_deg_, faster *r*_Act_to_Int_ (Decker and Hinton 2013; Roy and Singer 2015), slower translation initiation *r*_protein_syn_ (Hinnebusch 2011), and larger *K*_d_(0). We assume that the observed log-normal means of *r*_mRNA_deg_, *r*_protein_syn_, and *r*_Act_to_Int_ differ by 2-fold from the mean expected from mutational bias; for example, the mean of log_10_(*r*_mRNA_deg_) is −1.49, so the value of *μ* for *r*_mRNA_deg_ is – 1.49 + log_10_(2) = −1.19. We assume a larger bias for *K*_d_(0), namely that mutation is likely to reduce the affinity of a TF for a TFBS down to non-specific levels. Thus, we set *μ* = log_10_(*K*_d_(3)) = −5 for *K*_d_(0); note that in this case *μ* is equal to one of the boundary values, which will be hit far more often than during the evolution of other parameters. We assume that the observed central tendency estimate of protein stability does not depart from mutational equilibrium, therefore the value of *μ* for *r*_protein_deg_ is the mean of log_10_(*r*_protein_deg_) = −1.88.

The value of *σ* controls mutational effect size. We set the value of *σ* such that 1% of mutational changes from *x* = 10^*μ*^ go beyond the boundary values, for simplicity approximating by considering only the closer of the two boundary values on a log scale, i.e. we solve Supplementary Equation 17 for *σ*:

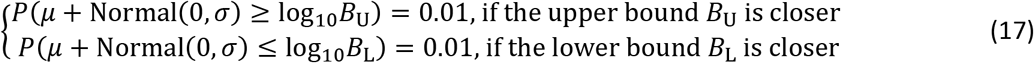

For example, the upper and the lower bounds of *r*_mRNA_deg_ are 0.54 min^−1^ and 7.5 × 10^−4^ min^−1^; on a log-scale, the upper bound is closer to 10^*μ*^ = 10^−1.19^ min^−1^. Plugging these values in Eq. S8 and solving for *σ*, we have *σ* = 0.396. We set the values of *σ* for *r*_protein_syn_, and *r*_protein_deg_ in the same way. However for *r*_Act_to_Int_, *σ* is set according to the lower bound, even though it is the more distant from 10^*μ*^, because otherwise a stable preinitiation complex will evolve too rarely. Under this high mutational variance, evolutionary outcomes at the two bounds are still only observed 5% of the time. For *K*_d_(0), because its upper bound is equal to 10^*μ*^, we set *σ* to 0.776, such that 1% of mutations can change the values of *K*_d_(0) by 100-fold or more.

Mutant values of *L*, *r*_Act_to_Int_, *r*_protein_syn_, *r*_protein_deg_, and *r*_mRNA_deg_ are constrained by the same bounds that constrain the initial values of these parameters (see previous sections). If a mutation increases the value of any of these 5 parameters to beyond the corresponding upper bound, we set the mutant value to the upper bound; similarly for a mutant value that is smaller than the lower bound of the corresponding parameter. For mutation to *K*_d_(0), we resample if *x*′ ≥ *K*_d_(3), because otherwise the mutation effectively “deletes” the TF by reducing its affinity to non-specific levels.

